# Directed evolution of a stem-helix–targeting antibody enables MERS-CoV cross-neutralization through enhanced binding affinity

**DOI:** 10.1101/2025.10.02.680065

**Authors:** Panpan Zhou, Meng Yuan, Yuexiu Zhang, Oliver Limbo, Ge Song, Fangzhu Zhao, Hejun Liu, Wan-ting He, Tazio Capozzola, Sean Callaghan, Gabriel Avillion, Xuduo Li, Nathan Beutler, Peter Yong, Fabio Anzanello, Thomas F. Rogers, Dennis R. Burton, Joseph G. Jardine, Ian A. Wilson, Raiees Andrabi

## Abstract

Broadly neutralizing antibodies (bnAbs) targeting conserved regions of the betacoronavirus spike are important for pan-betacoronavirus protection and pandemic preparedness. Here, we report on the isolation of a human monoclonal antibody, CC65.1, from a SARS-CoV-2 convalescent donor that targets the conserved S2 stem helix region. CC65.1 neutralizes various sarbecoviruses, including SARS-CoV-2, and binds the MERS-CoV spike but lacks MERS-CoV-neutralizing activity due to insufficient binding affinity. We utilized directed evolution to enhance the binding affinity of CC65.1 for the MERS-CoV S2 stem helix, yielding engineered antibody variants with newly acquired MERS-CoV-neutralizing activity. High-resolution structural analysis reveals key paratope mutations that optimize binding and stabilize epitope engagement. Our findings demonstrate the potential of rational antibody engineering to expand bnAb breadth across divergent betacoronaviruses. This work supports the development of engineered bnAbs and S2-targeted vaccines for broad betacoronavirus countermeasures and highlights strategies to achieve cross-lineage immunity for future pandemic threats.

**Author Summary:** The persistent emergence of new SARS-CoV-2 variants of concern that evade neutralizing antibody responses and other zoonotic betacoronaviruses with pandemic potential have provided strong motivation to develop broadly neutralizing antibodies (bnAbs) that target more conserved regions of the spike protein in sarbecoviruses and other betacoronaviruses. Here, we employed a directed evolution strategy to engineer the sarbecovirus-neutralizing antibody CC65.1, which targets the conserved S2 stem helix, to enhance its binding affinity with the MERS-CoV stem helix region, thereby conferring MERS-CoV neutralization. High-resolution structural studies of engineered CC65.1 revealed that key mutations reshape the paratope to better accommodate and stabilize the MERS-CoV S2 stem helix, resulting in increased binding affinity and neutralization potency. This study emphasizes the critical role of affinity maturation in expanding neutralization breadth and provides valuable insights for design of bnAbs to prevent and treat pandemic threats by betacoronaviruses.

## Introduction

The ongoing evolution and emergence of pathogenic coronaviruses, including Severe Acute Respiratory Syndrome Coronavirus 2 (SARS-CoV-2) and Middle East Respiratory Syndrome Coronavirus (MERS-CoV), continue to pose a significant threat to global public health [1–3]. The COVID-19 pandemic has underscored the urgent need for broadly protective countermeasures against current and future zoonotic coronaviruses [4, 5]. Among the four coronavirus genera, betacoronavirus harbors multiple high-risk pathogens, including SARS-CoV-1, SARS-CoV-2, and MERS-CoV, with the latter exhibiting a high case fatality rate and pandemic potential [6–9]. However, the antigenic diversity among betacoronavirus spike glycoproteins, particularly within the immunodominant receptor-binding domain (RBD), presents a formidable challenge for eliciting broadly neutralizing antibodies (bnAbs) that can cross-protect across betacoronavirus lineages [9–12]. The conserved S2 subunit of the spike protein, and specifically the stem helix region, has emerged as a promising target for cross-reactive antibody responses due to its structure and sequence conservation across betacoronavirus lineages [13–15].

Here, we report the isolation and characterization of a human monoclonal antibody (CC65.1) targeting the S2 stem helix that neutralizes SARS-CoV-2 and exhibits cross-reactivity with MERS-CoV spike, although does not neutralize MERS-CoV. To overcome this limitation, we utilized directed evolution to enhance binding of CC65.1 to the MERS-CoV stem helix region, thereby generating engineered antibody variants that gain MERS-CoV-neutralizing activity. High-resolution structural studies reveal the molecular basis of this cross-neutralization and provide mechanistic insights into epitope engagement and structural mimicry. We demonstrate how rational antibody engineering can be leveraged to broaden the neutralization breadth of existing antibodies toward distant betacoronavirus lineages, including MERS-CoV. Overall, our work provides a blueprint for designing pan-betacoronavirus bnAbs for intervention strategies and supports the development of broadly protective therapeutics and vaccines as critical components of preparedness against future pandemic coronaviruses as well as for seasonal betacoronaviruses.

## Results

### Isolation and characterization of the S2 stem helix bnAb CC65.1 from a SARS-CoV-2 convalescent human donor

To identify bnAbs targeting the conserved S2 stem helix of betacoronaviruses, we first screened sera from six SARS-CoV-2 convalescent donors (CC6, CC21, CC40, CC48, CC57, and CC65) for neutralizing activity against betacoronavirus pseudoviruses from different lineages, including SARS-CoV-2 (sarbecovirus) and MERS-CoV (merbecovirus) (Fig 1A-B). Among these, only CC40 and CC65 show cross-neutralization, with sera from donor CC65 exhibiting notably stronger neutralizing activity against MERS-CoV. Based on this observation, we focused on CC65 for the isolation of potential stem helix bnAbs. Except stem helix region, sequence identify between SARS-CoV-2 and HCoV-HKU1 (embecovirus) spike is low. Thus, spike proteins of SARS-CoV-2 and HCoV-HKU1 were selected as baits to sort CD3^-^CD4^-^CD8^-^CD14^-^CD19⁺CD20⁺IgG⁺IgM⁻ stem helix-specific B cells from peripheral blood mononuclear cells (PBMCs) of donor CC65 (Fig S1A). A total of 16 cross-reactive IgG⁺ B cells were isolated, and their paired heavy and light chain sequences were recovered and expressed as recombinant monoclonal antibodies (mAbs). These mAbs were screened for binding to the SARS-CoV-2 spike protein and to S2 stem helix peptides derived from SARS-CoV-1/2 and HCoV-HKU1 (Fig 1C-D). Eight of the 16 mAbs exhibit binding to the SARS-CoV-2 spike, and one (CC65.1) demonstrates the strongest stem helix binding activity (Fig 1D-E).

**Fig 1.**
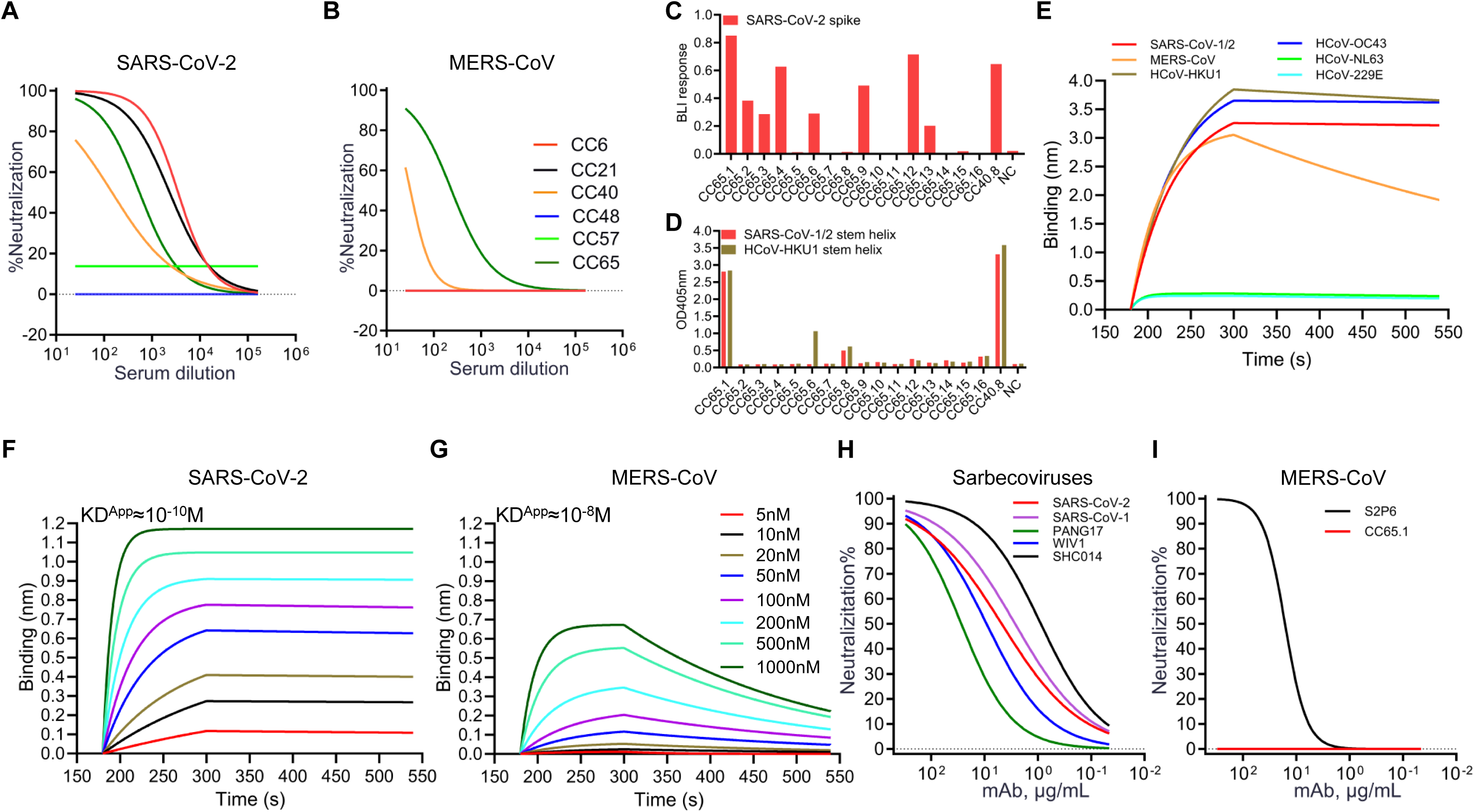
CC65.1 exhibits neutralizing activity against sarbecoviruses and broad binding across betacoronaviruses but does not neutralize MERS-CoV. (A-B) Neutralization curves of sera from SARS-CoV-2 recovered human donors (CC6, CC21, CC40, CC48, CC57, and CC65) to SARS-CoV-2 (A) and MERS-CoV (B). (C) BioLayer Interferometry (BLI) binding of 16 mAbs isolated from CC65 donor with SARS-CoV-2 recombinant soluble spike protein. (D) Binding of 16 isolated mAbs from CC65 with SARS-CoV-1/2 and HCoV-HKU1 S2 stem helix peptides by ELISA. The binding is shown as absorbance at OD_405_. S2 stem helix mAb, CC40.8, was used as a positive control [20] and supernatant from mock-transfected Expi293 cells as negative control. NC, negative control. (E) BLI binding curve of CC65.1 to S2 stem helix peptides from betacoronaviruses (SARS-CoV-1/2, MERS-CoV, HCoV-HKU1, and HCoV-OC43) and alphacoronaviruses (HCoV-NL63 and HCoV-229E). (F-G) BLI binding curves of CC65.1 to SARS-CoV-2 (F) and MERS-CoV (G) recombinant soluble spike proteins. CC65.1 was captured on a Protein A biosensor, followed by exposure to varying concentrations of spike protein. The KD^App^ was determined using a 1:1 binding kinetics model with ForteBio Data Analysis software. (H-I) Neutralization curves of CC65.1 against pseudoviruses of the clade 1a (SARS-CoV-1, WIV1, and SHC014), clade 1b (SARS-CoV-2 and Pang17) ACE2-utilizing sarbecoviruses (H) and MERS-CoV (I). The stem helix bnAb S2P6 was used as a positive control [18].

Based on the conserved nature of the S2 stem helix within betacoronaviruses, we next evaluated the breadth of CC65.1. The antibody binds both recombinant soluble and cell surface–expressed spikes of multiple betacoronaviruses, with stronger binding observed for SARS-CoV-1 and SARS-CoV-2 compared to MERS-CoV in both formats (Fig 1F-G and S1B). Consistent with its cell surface binding profile (Fig S1B), CC65.1 also binds S2 stem helix peptides from betacoronaviruses but not from the more phylogenetically distant alphacoronaviruses (HCoV- NL63 and HCoV-229E) (Fig 1E). We then assessed the neutralization breadth and potency of CC65.1 against a panel of sarbecoviruses, including clade 1a (SARS-CoV-1, WIV1, and SHC014) and clade 1b (SARS-CoV-2, Pang17), as well as MERS-CoV. CC65.1 neutralizes all tested sarbecoviruses with half-maximal inhibitory concentrations (IC₅₀) ranging from 0.87 to 26.97 µg/mL but fails to neutralize MERS-CoV, even at the highest tested concentration of 300 µg/mL (Fig 1H-I).

Given that neutralization potency often correlates with binding affinity [16, 17], we hypothesized that CC65.1 may exhibit insufficient binding affinity for the MERS-CoV spike or stem helix region to achieve neutralization. To test this notion, we compared the binding affinities of CC65.1 to various betacoronavirus recombinant soluble spikes and stem helix peptides. CC65.1 binds to SARS-CoV-1 and SARS-CoV-2 spikes and stem helix peptides with “apparent affinity” dissociation constants (KD^APP^) in the ∼10⁻¹⁰ M range (Fig 1F, S1C and Table S1). In contrast, binding to the MERS-CoV spike (KD^APP^≈10^-8^ M) and stem helix peptide (KD^APP^≈10^-9^ M) was approximately 100-fold and 10-fold weaker, respectively (Fig 1E, G and Table S1). Similar to other reported stem helix bnAbs [13], CC65.1 binds HCoV-HKU1 and HCoV-OC43 recombinant soluble spike proteins poorly (Fig S1D-E and Table S1). Together, these results demonstrate that CC65.1 targets the conserved S2 stem helix region, exhibiting broad binding across betacoronaviruses. However, its reduced binding affinity for MERS-CoV likely accounts for its lack of neutralizing activity against this virus.

### Mapping and characterization of CC65.1 binding hotspots on the SARS-CoV-2 and MERS-CoV stem helix peptides

The S2 stem region is conserved across betacoronavirus, including SARS-CoV-2, SARS-CoV-1, and MERS-CoV (Fig 2A). To understand the mechanism by which CC65.1 neutralizes sarbecoviruses but not MERS-CoV, we determined crystal structures of Fab CC65.1 with S2 stem helix peptides of SARS-CoV-1/2 and MERS-CoV at resolutions of 2.05 Å and 2.7 Å, respectively (Fig 2B and Table S2). In both structures, the S2 stem peptides display a helical conformation, with all six CC65.1 CDR loops involved in peptide binding (Fig 2B). CC65.1 belongs to the public class of antibodies encoded by IGHV1-46/IGKV3-20, represented by S2P6 [18] (Fig S2A-B). This antibody class also includes CC68.109, CC99.103 [13], COV89-22, COV30-14, and COV93-03 [19]. Compared to CC40.8, this class of antibodies binds the same S2 stem helix region [14, 20], but with a different binding angle and orientation (Fig S2B). The C-terminal region of the S2 stem helix is highly conserved between SARS-CoV-1/2 (AA1148-1156) and MERS-CoV (AA1231-1239) (Fig 2A). Both of these stem helix peptides insert into a hydrophobic groove formed by the heavy and light chains of CC65.1 (Fig 2B). For example, F^1148^ of SARS-CoV-1/2 and F^1231^ of MERS-CoV extensively stack with aromatic residues of CC65.1 including V_H_ W^100b^, F^97^ and V_L_ Y^91^, Y^32^, and F^96^. SARS-CoV-1/2 L^1152^ and Y^1155^ (corresponding to L^1235^ and F^1238^ in MERS-CoV) also form hydrophobic interactions with V_H_ H^35^, Y^33^, I^50^ and V_L_ F^96^, P^95a^ of CC65.1. Moreover, the conserved E^1151^ (SARS-CoV-1/2)/E^1234^ (MERS-CoV) forms a hydrogen bond with V_L_ Y^32^ (Fig 2C). In contrast, the N-terminal region of the epitope is not conserved between SARS-CoV-1/2 and MERS-CoV (Fig 2A). Residues ^1146^DS^1147^ of SARS-CoV-1/2 form three hydrogen bonds and salt bridges with CC65.1 V_L_ R^50^ and V_H_ W^100b^, while the corresponding ^1229^ID^1230^ in MERS-CoV do not form these interactions; instead I1229 forms hydrophobic interactions with V_L_ Y^32^ (Fig 2C). The loss of the polar interactions likely contributes to the reduced binding affinity of CC65.1 to MERS-CoV stem helix. Notably, V_L_ R^50^ is somatically mutated from its IGKV3-20 germline residue G^50^ that enables it to form these unique polar and charged interactions with SARS-CoV-1/2 (Fig S2C).

**Fig 2.**
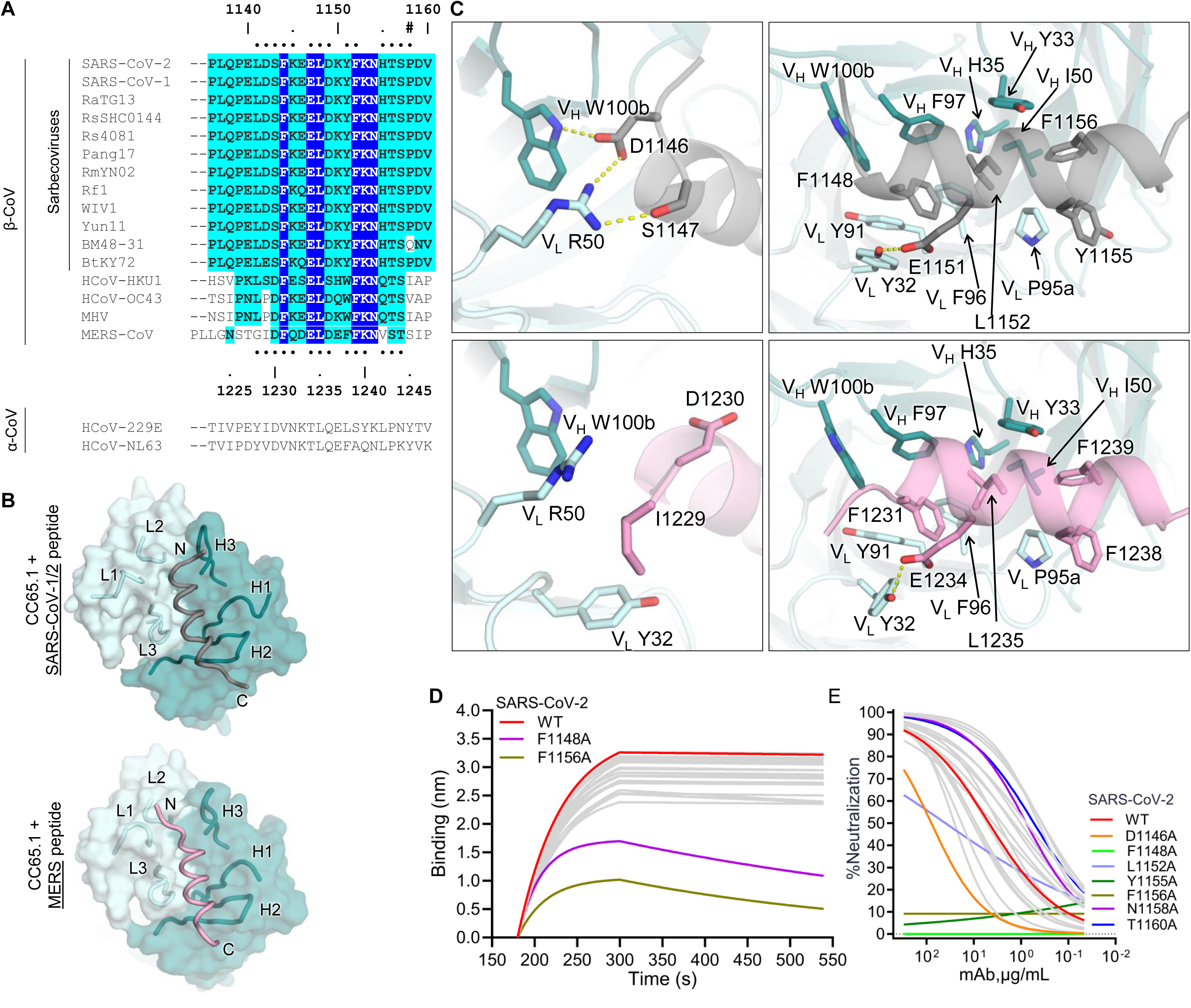
Crystal structures of CC65.1 in complex with SARS-CoV-1/2 and MERS-CoV S2 stem helix peptides. (**A**) Sequence alignment between the S2 stem helix region of betacoronaviruses. The SARS-CoV-2 numbering is shown on the top and MERS-CoV on the bottom. CC65.1 epitope residues (BSA > 0 Å^2^) of SARS-CoV-1/2 and MERS-CoV are indicated by dots on top and bottom of the alignment, respectively. Conserved identical residues are highlighted in blue, while similar residues are in cyan [amino acids that scored greater than or equal to 0 in the BLOSUM62 alignment score matrix were counted as similar here [38]. (**B**) Crystal structures of CC65.1 in complex with SARS-CoV-2 (grey) and MERS-CoV (pink) stem helix peptides. N- and C-terminus as well as CDR loops are indicated. Heavy and light chains of CC65.1 are shown in deep teal and pale cyan, respectively. (**C**) Details of molecular interactions between CC65.1 and SARS-CoV- 1/2 (top) and MERS-CoV (bottom) stem helix. S2 stem helix peptides of SARS-CoV-1/2 and MERS-CoV are shown in grey and pink, respectively. Hydrogen bonds and salt bridges are represented by yellow dashed lines. (**D**) BLI binding of CC65.1 to SARS-CoV-2 stem helix peptide alanine mutants spanning the whole peptide. The stem helix peptide mutants that most affect CC65.1 binding are shown in purple and olive in comparison to WT (red) and other stem helix mutants (grey). (**E**) Neutralization curves of CC65.1 to SARS-CoV-2 WT pseudovirus, pseudotyped variants with individual stem helix alanine mutants, and N^1158^ glycan-knockout SARS-CoV-2 pseudovirus variants (N1158A and T1160A).The WT virus is shown in red, and virus mutants that substantially affect CC65.1 neutralization and glycan-knockout mutants shown in assorted colors.

To confirm these epitopes recognized by CC65.1 and verify whether these residues are crucial for CC65.1 neutralization, we performed BLI binding and neutralization assays of CC65.1 to alanine scanning mutants of the SARS-CoV-2 stem helix epitopes and pseudoviruses, respectively. The F1148A and F1156A mutants substantially reduce binding of CC65.1 to SARS-CoV-2 stem helix peptide (Fig 2D and S2D). Mutations F1148A, Y1155A and F1156A, as well as D1146A and L1152A, reduce or abolish neutralization by CC65.1 (Fig 2E and S2D). As the SARS-CoV-2 has a glycosylation site at N^1158^, we generated N^1158^ glycan knockout mutants N1158A and T1160A to test the impact of N^1158^ on CC65.1 neutralization. Both mutations show ∼10-fold increased neutralization potency, indicating that glycosylation at N^1158^ has some impact on CC65.1 neutralization. Taken together, the S2 stem region is generally conserved among betacoronavirus, the interactions within the conserved amino acids leads to the recognition of SARS-CoV-2 and other betacoronaviruses spikes by CC65.1.

We then compared structures of six mAbs targeting the SARS-CoV-2 S2 stem helix region, including four mAbs isolated from COVID-19 convalescent patients: CC65.1 (this study), CC40.8 [20], S2P6 (another mAb that belongs to the IGHV1-46/IGKV3-20 class) [18], CV3-25 [21, 22], as well as two mAbs, B6 and IgG22, isolated from spike-immunized mice [23, 24] (Fig S2A-B). S2P6 binds with the same binding mode as CC65.1. Note that CC65.1 and S2P6 are both encoded by the same V genes, but with distinct HCDR3 loops (Fig S2C). Germline-encoded residues V_H_ Y^33^, H^35^, and V_L_ Y^32^, Y^91^of both mAbs form a hydrophobic groove that interacts with the hydrophobic core of the S2 stem helix (Fig S2C and S2E). A V_H_ S56G somatic hypermutation of CC65.1 avoids a possible clash between the side chain of V_H_ S56 and the antigen. Other IGHV1-46/IGKV3-20 antibodies, including CC68.109, CC99.103 [25], and COV89-22 [19], share the same mutation, demonstrating a common and convergent mutation upon affinity maturation. On the other hand, V_H_ S56 of S2P6 is mutated to a histidine, stacking with F1156 (Fig S2E).

Compared to CC65.1, the epitope of CC40.8 extends more toward the N-terminus of the S2 stem helix (Fig S2A), and interacts with a very different antibody binding mode (Fig S2B). Mouse antibodies B6 and Fab22 bind a more truncated epitope compared to CC65.1 that is translated along the groove between the heavy and light chains (Fig S2A-B). CV3-25 targets the S2 stem region with a completely different binding approach, and its epitope is more C-terminal compared to all the other mAbs analyzed here (Fig S2A-B) [20]. Finally, when modeled onto a SARS-CoV- 2 spike trimer in the prefusion state (Fig S2F), all of these S2 stem targeting mAbs that bind the hydrophobic face of the stem helix would clash with the adjacent protomers of the prefusion spike trimer [20].

### Directed evolution engineering of CC65.1 enables MERS-CoV neutralization through enhanced binding affinity

To further improve the neutralization potency of MERS-CoV as well as retain its broad reactivity, CC65.1 was affinity-matured using the MERS-CoV stem helix peptide. We employed the <u>S</u>ynthetic <u>A</u>ntibody <u>M</u>aturation by multiple <u>P</u>oint <u>L</u>oop library <u>E</u>n<u>R</u>ichments (SAMPLER), a directed evolution strategy that rapidly enhances the antibody affinity by systematically introducing mutations in the complementarity-determining regions (CDRs) [26, 27]. Briefly, separate heavy chain (HC) and light chain (LC) libraries were created by introducing single mutations into each of the CDR loops and displayed on the surface of yeast as molecular Fab (Fig 3A). The HC library was paired with an unmodified LC, and vice versa. These libraries underwent four rounds of selection to enrich variants exhibiting high affinity and specificity for the MERS-CoV stem helix peptide. In rounds one, two, and four, cells were labeled with subsaturating concentrations of biotinylated MERS-CoV stem helix peptide, and the top 5 to 10% of peptide-binding cells, normalized for Fab surface display, were enriched. In the third round, cells were incubated with biotinylated, detergent-solubilized Chinese hamster ovary cell membrane proteins (CHO-SMP) to remove polyreactive clones (Fig 3A). Subsequently, enriched HC and LC variants were recombined into a combinatorial HC/LC library and subjected to an additional four rounds of selection to identify optimal HC/LC pairs that exhibited maximal binding affinity to the MERS-CoV stem helix peptide.

**Fig 3.**
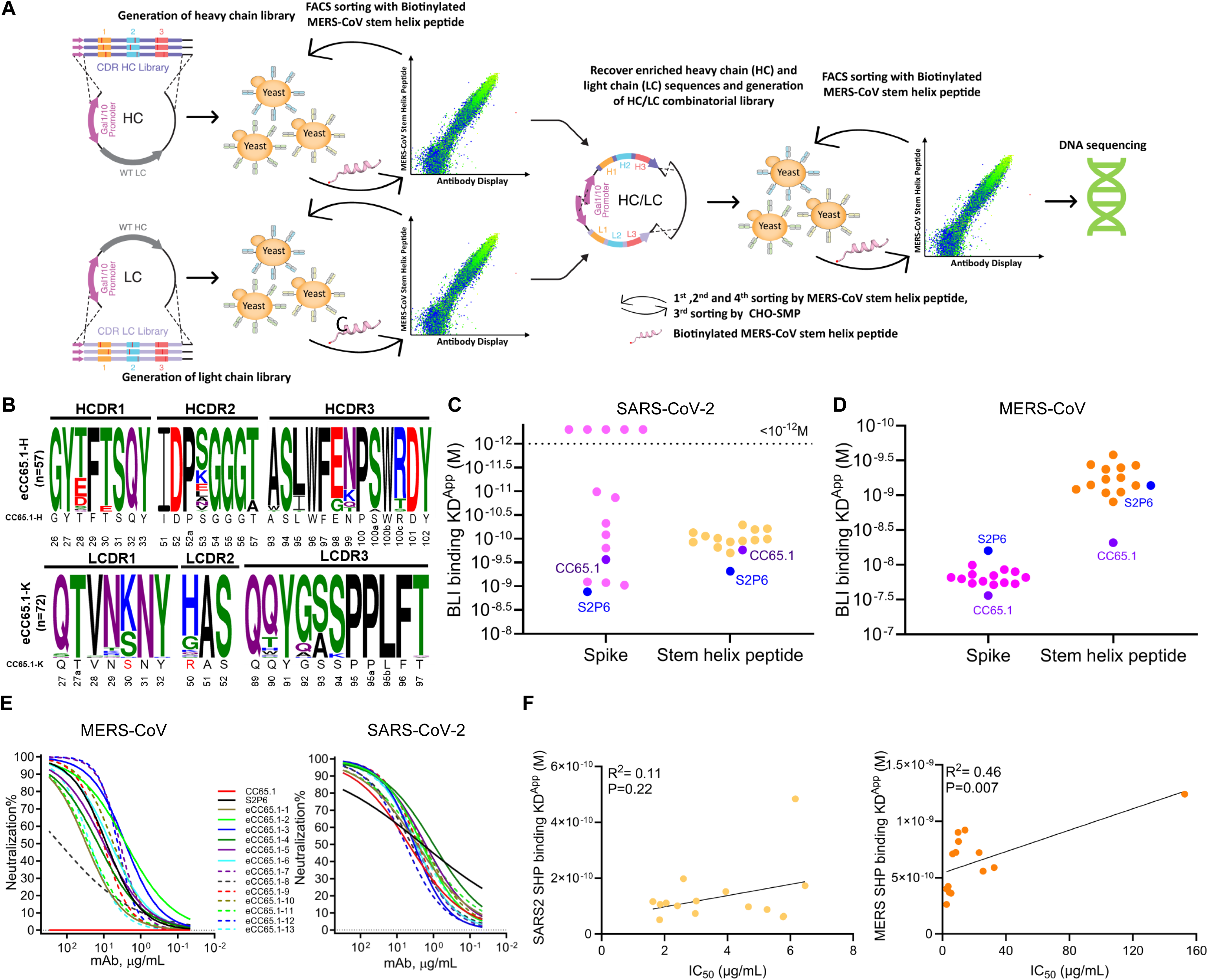
Directed evolution-guided engineering of the CC65.1 with enhanced binding affinity to the MERS-CoV stem helix peptide to enable it to neutralize MERS-CoV. (**A**) Library strategy: the CC65.1 antibody library with single mutations at each CDR loop was displayed as molecular Fab on the surface of yeast cells. The CC65.1 heavy chain (HC) library with up to three mutations was paired with the original light chain (LC), while LC library was paired with the original HC. After 4 rounds of FACS sorting by MERS-CoV stem helix peptide and CHO-SMP, the HC library and LC library were amplified and combined into combinatorial HC/LC library and further sorted for high binding clones by MERS-CoV stem helix peptide. Enriched clones with high binding affinities were sequenced, reformatted, and expressed as human IgG. (**B**) Identification of signature substitutions by comparison between CC65.1 and engineering CC65.1 (eCC65.1) variants using the program WebLogo. Clones from HC/LC library sort 4 were sequenced, sequences were aligned and sequence logo of CDRs of eCC65.1 HC and LC is shown. Sequences of CC65.1 CDRs are shown below each logo, where the signature substitutions enrich among the eCC65.1 antibodies are highlighted in red. Sequences are indicated by colors representing their different biochemical properties: green for polar, blue for basic, red for acidic, black for hydrophobic and purple for N or Q residues. Kabat numbering is shown under the sequence alignment. (**C-D**) Comparison of KD^App^ of S2P6, parental CC65.1 with selected eCC65.1 mAbs against recombinant soluble spike proteins and stem helix peptides from SARS-CoV-2 (C) and MERS-CoV (D). (**E**) Neutralization curves of parental CC65.1 and eCC65.1 mAbs against pseudotyped MERS-CoV (left) and SARS-CoV-2 (right) viruses. S2P6, a stem helix bnAb [18], was used as a positive control. (**F**) Plots showing correlation of the binding affinity (KD^APP^) of S2P6, CC65.1 and selected eCC65.1 mAbs to SARS-CoV-2 and MERS-CoV stem helix peptides with neutralization potency (IC_50_) against their corresponding pseudoviruses. Correlations were determined by nonparametric Spearman correlation two-tailed test with 95% confidence interval. The Spearman correlation coefficient (r squared) and p-values are indicated.

After the final selection round, compared with the cells from either HC or LC sort 4, the enriched variants of HC/LC sort 4 can bind the MERS-CoV stem helix peptide very strongly, even at very low peptide concentration (0.06 nM, HC/LC 92.4% vs HC 0.076% or LC 0.039%) (Fig S3). These variants also show cross-reactivity with SARS-CoV-1/2, HCoV-HKU1, and HCoV-OC43 stem helix peptides (Fig S4), confirming that the engineered CC65.1 (eCC65.1) mAbs retained broad binding activity. Plasmids encoding the HC and LC from more than 50 variants were amplified and sequenced. Sequence analysis of the engineered antibody variants from HC/LC sort 4 reveals mutations in CDRH1, CDRH2, and CDRH3 regions; however, none of these mutations shows strong enrichment (Fig 3B). In contrast, the light chain exhibits strong enrichment for two mutations: a serine-to-lysine substitution at position S30 in CDRL1 (S30K) and an arginine-to-histidine substitution at position R50 in CDRL2 (R50H) (Fig 3B). Compared with these two mutations, the enrichment of S93A in CDRL3 is slight.

A set of 13 eCC65.1 mAbs, named eCC65.1-1 through eCC65.1-13, were selected for further characterization (Fig 3C-F, S5 and S6A-C). To ensure that directed evolution did not inadvertently increase polyreactivity, we performed polyspecificity reagent (PSR) ELISA assay and also assessed polyreactivity or autoreactivity in HEp2 cells. All of the selected eCC65.1 mAbs are negative in the PSR assays and a few of selected eCC65.1 mAbs show some degree of polyreactivity or autoreactivity in HEp2 assay (Fig S5). All of them bind the MERS-CoV stem helix peptide with higher affinity than the parental CC65.1 antibody, showing an average 9.1-fold increase in KD^APP^ values (range: 3.9-18.4-fold) (Fig 3C and S6C). This affinity improvement is primarily driven by a reduced dissociation rate (Fig S6C). Binding to the MERS-CoV recombinant soluble spike protein also improved modestly (1.9-fold) compared to the parental CC65.1 (Fig 3C and S6C). Interestingly, most of the eCC65.1 variants exhibit a dramatic increase in binding affinity to the SARS-CoV-2 recombinant soluble spike protein (average 109-fold) and a moderate enhancement in binding to the SARS-CoV-2 stem helix peptide (1.91-fold), despite these targets not being part of the directed evolution selection process (Fig 3D and S6C). The disparity in binding affinity improvement between MERS-CoV and SARS-CoV-2 recombinant soluble spike proteins may reflect differences in epitope accessibility within the prefusion spike trimers of various betacoronaviruses.

To assess whether enhanced binding affinity translated into improved neutralization potency, IC_50_ values against MERS-CoV, SARS-CoV-2, and SARS-CoV-1 were determined (Fig 3E and S6C). eCC65.1 variants display the similar neutralization potency against SARS-CoV-2 and SARS-CoV- 1 with the parental antibody. However, all engineered variants, including eCC65.1-6 (V_L_ S30K and R50H), eCC65.1-12 (V_L_ S30K R50H, and S93A) and eCC65.1-13 (V_L_ S30K, R50G and S93A), which only have mutations in light chain, show significantly improved MERS-CoV neutralization, with a median IC_50_ of 22.3 µg/mL (Fig 3E and S6A-C). eCC65.1-6 and eCC65.1-12 have higher MERS-CoV neutralization potency than eCC65.1-13, and eCC65.1-1, 4, 5, 8, 9, 11 and 13, which are without V_L_ S30K or R50H mutation, are not as potent as the others for MERS-CoV neutralization. Thus, these two mutations may have a greater effect on MERS-CoV neutralization potency than the other mutations. For naturally derived KV3-20 MERS-CoV-neutralizing human stem helix bnAbs, although basic amino acids (K and R) enriched on CDRL1 site 30 (7/22), V_L_ K30 (2/22) and H50 (0/22) seem to be rare among these antibodies,and all eight KV3-20 MERS-CoV-non-neutralizing antibodies lack both V_L_ K30 and H50 (Fig S6D). In order to assess the role of these two mutations, as well as S93A which is enriched slightly in CDRL3, in improving MERS-CoV neutralization potency, eCC65.1-12 was selected for structural study.

The neutralization potency of eCC65.1 mAbs reveals a positive correlation with binding affinity to the MERS-CoV stem helix peptide, but not the MERS-CoV recombinant soluble spike protein (Fig 3F and S6E). For SARS-CoV-2, binding affinity to both the stem helix peptide and recombinant soluble spike protein show a positive correlation with neutralization potency, although the correlation with spike protein is not significant (Fig 3F and S6E) .Despite a >100-fold increase in binding affinity to the SARS-CoV-2 recombinant soluble spike protein for most of the eCC65.1 mAbs, no substantial improvement in SARS-CoV-2 neutralization potency is observed (Fig 3E and S6C). This may be due to the parental CC65.1 already exhibiting SARS-CoV-2 neutralization (IC_50_ = 6.16 µg/mL), suggesting that further binding affinity enhancement does not translate into improved neutralization potency beyond a certain threshold. In contrast, because CC65.1 lacks baseline MERS-CoV neutralizing activity, binding affinity improvements in eCC65.1 variants lead to a marked gain in neutralization capability [28]. Together, these results demonstrate that using the SAMPLER platform for directed evolution, we successfully engineered CC65.1 variants with enhanced binding affinity and newly acquired MERS-CoV-neutralizing activity. This highlights the potential of affinity maturation strategies to expand antibody breadth and functionality against related, previously resistant viruses.

### Structural basis of MERS-CoV neutralization by eCC65.1

To further understand the mechanism by which enhanced binding affinity to the MERS-CoV stem helix peptide contributes to neutralization potency against MERS-CoV virus, we selected eCC65.1-12, one of the mAbs with improved binding and neutralization activity against MERS-CoV that contains only three mutations in light chain (V_L_ S30K, R50H, and S93A) for detailed functional characterization (Fig 4). We compared the neutralization potency and breadth of parental CC65.1 and eCC65.1-12 against sarbecoviruses (SARS-CoV-1, SARS-CoV-2, SHC014, PANG17, and WIV1) and MERS-CoV. Both mAbs effectively neutralize all sarbecoviruses tested and eCC65.1-12 neutralizes MERS-CoV with an IC_50_ of 4.18 µg/mL (Fig S7A). Additionally, eCC65.1-12 retains the broad-reactivity of parental CC65.1, effectively binding to stem helix peptides and cell surface-expressed spike proteins from betacoronaviruses, but not alphacoronaviruses, such as HCoV-NL63 and HCoV-229E (Fig S7B-C, and Table S1). As the stem helix is conserved in betacoronaviruses, the improved binding affinity to MERS-CoV stem helix contributes to the substantially increased binding affinity to stem helix of other betacoronaviruses, such as SARS-CoV-1/2 (35.4-fold), HCoV-HKU1(6.7-fold) and HCoV-OC43 (7.6-fold), but doesn’t significantly improve the binding affinity to recombinant soluble spike proteins except SARS-CoV-2 (Fig 1F-G, S1C-E, S7D-H and Table S1). However, the binding affinity of eCC65.1-12 to cell surface-expressed full-length HCoV-HKU1 and MERS-CoV spike are enhanced (Fig S1B, S7C and Table S1). Overall, eCC65.1-12 is endowed with broad betacoronavirus cross-reactivity and sarbecovirus and MERS-CoV neutralizing activity.

**Fig 4.**
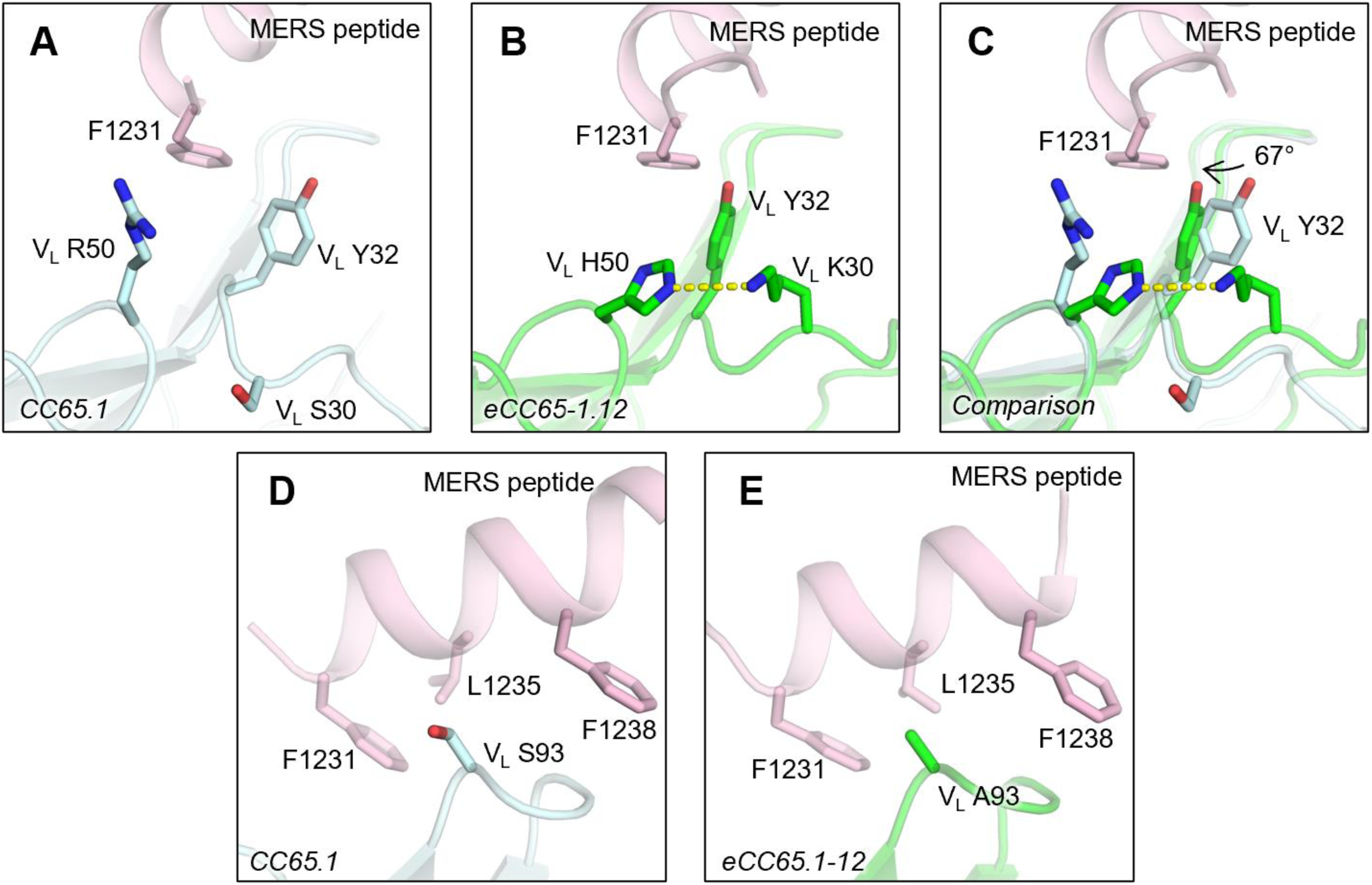
Structural basis of eCC65.1-12 with improved binding affinity against the S2 stem helix of MERS-CoV. Interaction of the MERS-CoV S2 stem helix peptide with CC65.1 and eCC65.1-12 from their crystal structures. CC65.1 is shown in pale cyan and eCC65.1-12 in green, and MERS-CoV peptide is in pink. Hydrogen bonds are represented by yellow dashed lines. The effect of mutations compared to WT in eCC65.1-12 at V_L_ residues 30 and 50 are shown in panels A-C and at V_L_ residue 93 in panels D and E.

We further determined a crystal structure of eCC65.1-12 in complex with the MERS-CoV S2 stem helix peptide (Fig 4). Compared with parental CC65.1, the eCC65.1-12 contains three mutations in the LC, including S30K, R50H, and S93A. The R50H substitution induces a 67° rotation in the rotamer of the V_L_ Y^32^ side chain that enables V_L_ Y^32^ of eCC65.1-12 to form a T-shaped stacking interaction with MERS-CoV F1231 which is higly conserved among betacoronaviruses (equivalent to SARS-CoV-2 F1144) (Fig 2A), likely strengthening the interaction (Fig 4A-C). Indeed, in most of the eCC65.1 mAbs, V_L_ R^50^ is mutated, mainly to histidine (Fig 3B and S6B). Interestingly, eCC65.1-13 is almost identical to eCC65.1-12 except for one residue difference at V_L_ residue 50 (Fig S6B), where eCC65.1-13 contains a glycine compared to histidine in eCC65.1-12. This one-residue difference leads to a ∼6-fold difference in neutralization IC_50_ (Fig S6C), which further suggests enhanced binding due to the V_L_ H^50^ mutation. V_L_ S30K now forms a hydrogen bond with V_L_ H^50^ that may further stabilize the CDR conformations due to these new interactions between CDR L1 (K30, Y32) and L2 (H50) (Fig 4B-C).V_L_ S93A also increases hydrophobic interactions with epitope residues F1231, L1235, and F1238 in MERS-CoV (Fig 4D-E).

## Discussion

The emergence of pathogenic betacoronaviruses, such as SARS-CoV-1, SARS-CoV-2 and MERS-CoV, underscore the urgent need for bnAbs that can provide cross-lineage protection [29, 30]. In this study, we report the isolation of a human mAb, CC65.1, from a SARS-CoV-2 convalescent donor that targets the conserved S2 stem helix region of the spike protein. CC65.1 neutralizes SARS-CoV-2 and other ACE2-utilizing sarbecoviruses and exhibits cross-reactive binding to MERS-CoV, although without neutralizing activity. Structural and biochemical analyses reveals that the lack of MERS-CoV neutralization is attributable to reduced binding affinity, driven in part by sequence divergence at the N-terminal end of the S2 stem helix. To overcome this, we employed a directed evolution strategy using the SAMPLER platform to enhance the binding affinity of CC65.1 to the MERS-CoV stem helix region. The resulting engineered variants, for example eCC65.1-12, acquire MERS-CoV-neutralizing activity while retaining broad reactivity to other betacoronaviruses. High-resolution structural studies of eCC65.1-2 reveals the mechanistic basis for MERS-CoV neutralization potency improvenment through affinity maturation, highlighting key light chain mutations, such as S30K in CDRL1 and R50H in CDRL2, which reshape the paratope to better accommodate and stabilize the MERS-CoV S2 stem helix. The parental antibody, CC65.1, forms extensive interactions with the N-terminal two amino acids ‘^1146^DS^1147^’ of SARS-CoV-2 stem helix, which differs from those in MERS-CoV ‘^1229^ID^1230^’. Our analysis also provides structural explanation of how the matured antibody, eCC65.1-2, has mutated its paratope to make stronger interactions with MERS-CoV S2 stem helix. Similar to this study, we previously analyzed how an in vitro affinity-matured mAb targeting the conserved CR3022 site of the SARS-CoV-2 RBD gained neutralization potency and breadth against SARS-CoV-2 variants while retaining SARS-CoV-1 neutralization [26]. Taken together, these findings underscore how even subtle epitope differences within a conserved region can significantly affect neutralization and how rational engineering can overcome these barriers.

Our study adds to the growing body of evidence supporting the S2 stem helix as a viable target for broad betacoronavirus interventions [18, 31–33]. While the receptor-binding domain (RBD) remains the dominant target of most neutralizing antibodies, its high variability across and within betacoronavirus lineages limits its utility for pan-betacoronavirus intervention strategies [34, 35]. In contrast, the S2 stem helix is conserved in both structure and sequence and has critical function in virus entry,which makes it an ideal target for broad-spectrum therapeutics [13, 19, 20, 36]. However, as our work demonstrates, some natural antibodies like CC65.1 may require binding affinity optimization to neutralize more antigenically distinct viruses such as MERS-CoV. This work demonstrates that rational antibody engineering—guided by structural insights and high-throughput selection—can expand the breadth of existing bnAbs and enable cross-lineage neutralization, including against viruses not originally targeted by the immune response. Such approaches could be readily applied to optimize existing antibodies from virus-infected people or vaccine-induced antibodies to cover the full diversity of pathogenic and pre-emergent betacoronaviruses.

In conclusion, the successful engineering of CC65.1 into a cross-neutralizing antibody against both sarbecoviruses and MERS-CoV provides a blueprint for development of next-generation countermeasures. Future efforts should focus on combining broad S2-targeting antibodies with complementary RBD-targeting bnAbs or incorporating engineered stem helix immunogens into vaccine platforms [37]. These strategies will be vital for establishing broad and durable immunity against both current and future betacoronavirus threats and for advancing pandemic preparedness.

## Materials and methods

### Cell lines

FreeStyle293-F cells (Thermo Fisher Scientific Cat# R79007) were cultured in FreeStyle 293 Expression Medium (Gibco Cat# 12338018), and Expi293F cells (Gibco Cat# A14527) were maintained in Expi293 Expression Medium (Gibco Cat# A1435101). Both suspension cells were incubated in the shaker at 150 rpm, 37°C, 8% CO_2_. HEK293T cells, Hela cells stably expressing hACE2 (Hela-hACE2) and hDPP4 (Hela-hDPP4) cells were grown in Dulbecco’s Modified Eagle Medium (DMEM) supplemented with 10% heat-inactivated FBS, 4mM L-Glutamine and 1% penicillin-streptomycin, in an incubator at 37°C, 5% CO_2_.

### Expression and purification of betacoronavirus spike protein and antibody

The betacoronavirus recombinant soluble spike proteins with a C-terminal His-Avi-tag were expressed in FreeStyle293-F cells as previously described [39]. For purification, cell culture supernatants were collected and secreted spike proteins were purified with the HisPur Ni-NTA Resin (Thermo Fisher Scientific Cat# 88221) and eluted with 200 mM imidazole. The purified proteins were concentrated and further purified by size-exclusion chromatography by Superdex 200 Increase 10/300 GL column (GE Healthcare Cat# GE28990944) in PBS. Proteins were concentrated again and stored at -80°C for further use.

Monoclonal antibody (mAb) expression and purification were performed as previously reported [39]. In brief, the paired heavy and light chain were co-transfected into Expi293 cells using FectoPRO PolyPlus reagent (Polyplus Cat# 116–040). After 24h post-transfection, sodium valproic acid and glucose were added. After 4 days of incubation, cell supernatants were harvested, and mAbs were purified using Protein A and Protein G Sepharose (GE Healthcare Cat# 17061805).

### Flow cytometry B cell profiling and monoclonal antibody isolation

To isolate antigen-specific memory B cells, HCoV-HKU1 and SARS-CoV-2 spike were used as probes for single cell sorting. B cell sorting and antibody isolation were performed as previously described [39]. The frozen PBMCs from donor CC65 were thawed and recovered in 10mL RPMI 1640 medium containing 50% FBS immediately before staining. The following reagents were used during staining: CD3 (APC Cy7, BD Pharmingen Cat# 557757), CD4 (APC-Cy7, Biolegend, Cat# 317418), CD8 (APC-Cy7, BD Pharmingen Cat# 557760), CD14 (APC-H7, BD Pharmingen Cat# 561384), CD19 (PerCP-Cy5.5, Biolegend Cat# 302230), CD20 (PerCP-Cy5.5, Biolegend Cat# 302326), IgG (BV786, BD Horizon Cat# 564230) and IgM (PE, Biolegend Cat# 314508). After staining by the above Ab mixture, cells were incubated with HCoV-HKU1 spike and SARS-CoV-2 spike which were conjugated to streptavidin-AF488 (Thermo Fisher Scientific Cat# S11223) and streptavidin-AF647 (Thermo Fisher Scientific Cat# S21374), respectively, on ice for 30min. Prior to sorting, FVS510 Live/Dead stain (Thermo Fisher Scientific Cat# L34966) was added to exclude the dead cells. Cross-reactive Spike-protein specific B cells (SARS-CoV-2^+^HCoV- HKU1^+^CD19^+^CD20^+^CD3^-^CD4^-^CD8^-^CD14^-^IgM^-^IgG^+^) were sorted into 96-well plates. To amplify the variable regions of lgG heavy and light chain, RT-PCR and nested PCR were performed as previously described [13]. The purified DNA fragments were cloned into expression vectors encoding human IgG1, and Ig kappa/lambda constant domains, respectively, using HiFi DNA assembly (New England Biolabs Cat# E2621L).

### ELISA

The binding of 16 isolated mAbs from CC65 donor with SARS-CoV-1/2 and HCoV-HKU1 S2 stem helix peptides was assessed by ELISA. The 96-well half-area microplates were coated with 100 ng/well streptavidin (Jackson Immuno Research Labs Cat# 016-000-084) at 4°C overnight. The plates were washed three times with PBST (PBS + 0.05% Tween20) and blocked with 3% bovine serum albumin (BSA) in PBS for 2h at room temperature (RT). After removing the blocking buffer, plates were treated with biotinylated S2 stem helix peptides (5 μg/mL in 50 μL 1% BSA) for 1h at RT. Following additional washes, diluted antibodies (10 μg/mL in 50 μL 1% BSA) were added and incubated for 1h. Following washes, secondary antibody (Jackson ImmunoResearch Laboratories Cat# 109-055-008) was added in 1:1000 dilution for an additional 1h. After final washes, alkaline phosphatase substrate (Sigma-Aldrich Cat# S0942-200TAB) was added. Absorbance at 405 nm was measured after 30 min using VersaMax microplate reader (Molecular Devices).

### Antibody library generation

The CC65.1 heavy chain (HC) and light chain (LC) Fab libraries were generated as reported previously [26, 28, 40]. In brief, oligopools, which contained a single mutation in each complementarity-determining region (CDR), were synthesized (Integrated DNA Technologies). Then, the CDR1/2/3 mini-libraries were assembled into combinatorial heavy chain and light chain libraries. The pYDSI2w vector containing the bidirectional Gal1-10 promoter was used to display the Fab libraries on the yeast surface. V5 and c-Myc epitope tag at the C-terminal of HC and LC, respectively, were taken to measure the amount of Fab displayed on the yeast surface. The HC library was generated by cloning the HC CDR1/2/3 library into pYDSI2w vector, which already had the parental CC65.1 LC. The LC library was generated by cloning the LC CDR1/2/3 library into pYDSI2w vector, which already had the parental CC65.1 HC. In order to generate CC65.1 HC/LC combinational library, the HC and LC sequences were amplified from the sorted HC and LC libraries with primers overlapping in the Gal1-10 promoter. Then, the purified HC and LC fragments were ligated by HiFi DNA assembly (New England Biolabs Cat# E2621L) and amplified to get the LC–Gal1-10–HC product which could be inserted into the empty pYDSI2w vector to generate CC65.1 HC/LC library.

### Yeast transformation

The *Saccharomyces cerevisiae* YVH10 electrocompetent cells were mixed with 1 μg linearized pYDSI2w vector and 5 μg HC, LC or HC/LC DNA, then were transferred into a 0.2 cm electroporation cuvette (Bio-Rad Cat# 1652086) and inserted into a Gene Pulser Xcell Electroporation System (Bio-Rad Cat# 1652666) using following settings: square wave, voltage = 500 V, pulse length = 15.0 ms, number of pulses = 1, pulse interval = 0, and cuvette = 2 mm. After electroporation, the yeast cells were moved into 25 mL YPD medium and cultured at 30°C for 1 hour with shaking at 200 rpm. Then, 2.5 μL of the yeast cells was diluted serially to estimate transformation efficiency with the colony-forming unit assay on synthetic complete agar plates without tryptophan (SC-Trp) (Sunrise Science Cat# 1710-300). The other cells were transferred to 250 mL SC-Trp medium (Sunrise Science Cat# 1709-500) with 1% penicillin/streptomycin (Corning Cat# 15323671) and shaken overnight at 30°C for further sorting.

### Yeast library labeling and sorting

After yeast transformation, the cells were passaged 1:20 next day, then induced at OD=1.0 overnight in SGCAA medium [41]. For each library, 5 x 10^7^ cells were stained in the first round of sorting, 1 × 10^7^ cells in the following other 3 rounds. After being spun down and washed by PBSA (PBS with 1% BSA), the cells were incubated with biotinylated stem helix peptides or detergent-solubilized Chinese hamster ovary cell membrane proteins (CHO-SMP) at several non-depleting concentrations respectively for 30 min at 4°C. After washing by PBSA, the yeast cells were stained by anti-c-Myc antibody (FITC, Immunology Consultants Laboratory Cat# CMYC-45F), anti-V5 antibody (AF405, made in house), and streptavidin-APC (Invitrogen Cat# CSA1005) in 1:100 dilution for 20 min at 4°C. After final washing, the cells were resuspended in 1 mL PBSA and loaded on BD FACSMelody cell sorter, and top 5-10% of cells with high binding activity to a certain stem helix peptide concentration were sorted. Sorted cells were cultured in 2 mL SC-Trp medium (Sunrise Science Cat# 1709-500) supplemented with 1% Penicillin/Streptomycin (Corning Cat# 15323671) at 30°C overnight for further use.

### Yeast colony and DNA sequencing

After culturing the cells from HC/LC sort4 in SC-Trp medium (Sunrise Science Cat# 1709- 500)overnight, the cells were diluted serially and grown on SC-Trp plates at 30°C overnight, then single colonies were picked and grown in SC-Trp medium at 30°C overnight. The DNA was extracted from yeast cells as reported previously [26, 28, 40], and the heavy chain and light chain regions were amplified from the DNA and Sanger sequenced. Bioedit (https://bioedit.software.informer.com) and WebLogo (https://weblogo.berkeley.edu/) were used to analyze the sequences.

### Pseudovirus production and neutralization assay

Sarbecovirus and MERS-CoV pseudoviruses were generated in HEK293T cells. Briefly, 12.5 μg pCMV-dR8.2 dvpr (Addgene Cat# 8455), 10 μg pBOB-Luciferase (Addgene Cat# 170674), and 2.5 μg sarbecovirus or MERS spike plasmid were mixed with transfection reagent Lipofectamine 2000 (ThermoFisher Scientific Cat# 11668019) and incubated for 15 min at RT and then transferred into HEK 293T cells. After 12-16h, the medium was changed with fresh complete medium (10% FBS, 4 mM L-Glutamine, and 1% Penicillin/Streptomycin). Supernatants containing pseudovirus were harvested after 48h post transfection, then aliquoted and frozen at -80 °C for further use.

Neutralization assay for ACE2-utilizing sarbecoviruses including clade 1a (SARS-CoV-1, WIV1, SHC014) and clade 1b (SARS-CoV-2, Pang17) was performed by HeLa-hACE2 cell, MERS-CoV neutralization was performed using Hela-hDPP4 cells. The 3-fold serially diluted antibodies were incubated with the same volume (25 μL/well) of pseudovirus for 1h at 37°C. After incubation, 50 μL of Hela-hACE2 or Hela-hDPP4 cells (10,000 cells/well) were added to each well. After 48h of incubation, luciferase activity was measured by BrightGlo substrate (Promega Cat# E2620) according to the manufacturer’s instructions. Fifty percent maximal inhibitory concentrations (IC_50_s) were determined by the dose-response-inhibition model with 5-parameter Hill slope equation in GraphPad Prism 7 (GraphPad Software).

### HEp2 epithelial cell polyreactive assay

Polyreactivity of engineered CC65.1 antibodies to human epithelial type 2 (HEp2) was determined by indirect immunofluorescence using HEp2 slides (Hemagen Cat# 902360). Briefly, mAb was diluted to 50 μg/mL in PBS and added onto immobilized HEp2 slides, followed by incubation for 30 min at RT. Slides were washed three times with PBS, and one drop of FITC-conjugated goat anti-human IgG was added onto each well and incubated for 30 min in the dark at RT. After washing, the coverslip was added to HEp2 slide with glycerol and the images were photographed on a Nikon fluorescence microscope to detect FITC signal.

### Polyspecificity reagent (PSR) ELISA

Solubilized CHO cell membrane protein (SMP), human insulin (Sigma-Aldrich Cat# I2643), single strand DNA (Sigma-Aldrich Cat# D8899) were coated onto 96-well half-area high-binding plates (Corning Cat# 3690) at 5 μg /mL in PBS and incubated overnight at 4°C. After washing with PBST, plates were blocked with 3% BSA for 2h at 37°C. The 5-fold serially diluted antibody starting from 50 μg/mL was added in plates to incubate for 1h at RT. The assay was performed as described in section ‘‘ELISA’’.

### CELISA binding

Binding of CC65.1 and eCC65.1-12 to cell surface expressed spikes from different coronaviruses was evaluated by cell-based ELISA (CELISA) as described previously [25]. A total of 4x10^6^ HEK293T cells were seeded into each 10cm culture dish and incubated at 37°C. After 24h, HEK293T cells were transfected with plasmids encoding full-length coronavirus spikes. After incubation at 37°C for another 48 hours, the cells were harvested and distributed into each well of 96-well round-bottom tissue culture plates (Corning Cat# 3799) for individual staining. Before staining, cells were washed three times with 200 µL FACS buffer (1xPBS, 2%FBS, 1mM EDTA), then were stained for 1h on ice with CC65.1 or eCC65.1-12 at 10 μg /mL in 50 µL staining buffer. After washing another three times with FACS buffer, the cells in each well were stained with 50 µL FACS buffer containing anti-human IgG Fc antibody (PE, diluted at 1:200, Southern Biotech Cat# 904009) and Zombie-NIR viability dye (diluted at 1:1000, BioLegend Cat# 423105) on ice in dark for 45min. Following the final washes with FACS buffer, the cells were resuspended and analyzed by flow cytometry (BD Lyrics cytometer), and the binding data were generated by calculating the Mean Fluorescence Intensity (MFI) using FlowJo 10 software. Mock-transfected 293T served as a negative control.

### BioLayer Interferometry binding (BLI)

The binding of parental Ab CC65.1 and engineered CC65.1 with betacoronavirus spike proteins and stem helix peptides was determined by BLI using the Octet RED96e system. For binding to spike protein, antibody was captured by anti-human IgG Fc capture (AHC) biosensors (ForteBio Cat# 185063) for 60s, followed by 60s baseline step with Octet buffer to remove unbound antibody. The sensors were then immersed into the spike protein in Octet buffer (PBS with 0.1% Tween) for 120s for association, followed by transferring into Octet buffer for 240s for dissociation. For stem helix peptide binding, N-terminal biotinylated stem helix peptides were diluted in Octet buffer and captured by the streptavidin (SA) biosensors (ForteBio Cat# 185020) for 60s, then the sensors were transferred into Octet buffer for 60s to remove the unbound peptides, followed by associating with monoclonal antibodies for 120s and dissociating in Octet buffer for 240s. The data were analyzed using the ForteBio Data Analysis software for correction, and the kinetic curves were fit to a 1:1 binding mode.

### Expression and purification of Fab and single-chain variable fragment (scFv)

CC65.1 and eCC65.1-12 Fab plasmids were generated by introducing the stop codon right after the amino acid “KSC” in the heavy chain constant region. The truncated heavy chains were co- transfected with the corresponding light chains in Expi293F cells, and the supernatants were harvested 4 days post transfection. Fabs were purified with CaptureSelect™ CH1-XL Affinity Matrix (Thermo Fisher Scientific Cat# 1943462250) and Superdex 200 Increase 10/300 GL column (GE Healthcare Cat# GE28-9909-44). The CC65.1 scFv genes were cloned with a C- terminal His_6_-tag in the heavy chain and light chain variable region (VH-VL) orientation, which linked together by a (G4S)_3_ to form a VH-VL-(G4S)_3_-His_6_ format. The production and purification methods of scFv with His tag are very similar with those in “Expression and purification of HCoV S-proteins and antibody” section.

### Crystallization and X-ray structure determination

A mixture of antibodies and 10 × (molar ratio) stem helix peptides were screened for crystallization using the 384 conditions of the JCSG Core Suite (Qiagen) on our robotic CrystalMation system (Rigaku) at Scripps Research. Crystallization trials were set-up using the vapor diffusion method in sitting drops containing 0.1 μL of protein and 0.1 μL of reservoir solution. Diffraction-quality crystals were obtained in the following conditions: CC65.1 scFv / SARS-CoV-2 S2 stem peptide (11 mg/mL): 20% (w/v) PEG3350 and 0.2 M CaCl_2_ at 20°C; CC65.1 Fab / MERS-CoV S2 stem peptide (15 mg/mL): 2% (v/v) PEG400, 2 M ammonium sulfate, and 0.1 M HEPES pH 7.5 at 20°C; eCC65.1-12 Fab / MERS-CoV S2 stem peptide (16 mg/mL): 0.2 M ammonium sulfate, 25% (w/v) PEG 4000, 0.1 M sodium acetate pH 4.6 at 20°C

All crystals appeared on day 7 and were harvested on day 10. Before flash cooling in liquid nitrogen for X-ray diffraction studies, crystals were equilibrated in reservoir solution supplemented the following cryoprotectants: CC65.1 scFv / SARS-CoV-2 S2 stem peptide: 20% ethylene glycol; CC65.1 Fab / MERS-CoV S2 stem peptide: 20% ethylene glycol; eCC65.1-12 Fab / MERS-CoV S2 stem peptide: 10% ethylene glycol.

Diffraction data were collected at cryogenic temperature (100 K) at the Advanced Light Source on beamline 5.0.1, Stanford Synchrotron Radiation Lightsource (SSRL) on Scripps/Stanford beamline 12-1, and Advanced Photon Source at the Argonne National Laboratory on beamline 23-ID-B. Data were processed with HKL2000 [42] (Table S2). Structures were solved by molecular replacement (MR) using PHASER [43] with PDB 7SJS as the MR model. Iterative model building and refinement were carried out in COOT and PHENIX [44, 45], respectively (Table S2). Epitope and paratope residues, as well as their interactions, were identified by accessing PISA at the European Bioinformatics Institute (http://www.ebi.ac.uk/pdbe/prot_int/pistart.html) [43].

### Statistical Analysis

Statistical analysis was performed using Graph Pad Prism 7, USA. IC_50_ titers were compared using the non-parametric unpaired Mann-Whitney-U test. Correlations were determined by nonparametric Spearman correlation two-tailed test with 95% confidence interval. The Spearman correlation coefficient (r squared) and p-value are indicated. Groups of data were compared using the Kruskal-Wallis non-parametric test. Dunnett’s multiple comparisons test were also performed between experimental groups. Data were considered statistically significant when p < 0.05.

## Acknowledgements

This work was supported by National Institutes of Health-(NIH), National Institute of Allergy and Infectious Diseases-(NIAID) awards, R01 AI170928 (R.A.), R01 AI190286 (M.Y., D.R.B, I.A.W.) and the Gates Foundation INV-004923 (I.A.W., D.R.B.). We thank Henry Tien for technical support with the crystallization robot. We are grateful to the staff of the Advanced Light Source beamline 5.0.1, Advanced Photon Source beamline 23-ID-B, and Stanford Synchrotron Radiation Lightsource (SSRL) beamline 12-1 for assistance. GM/CA@APS has been funded by the National Cancer Institute (ACB-12002) and the National Institute of General Medical Sciences (AGM-12006, P30GM138396). This research used resources of: the Advanced Light Source, a U.S. DOE Office of Science User Facility under contract no. DE-AC02-05CH11231, the Advanced Photon Source; a U.S. Department of Energy (DOE) Office of Science User Facility operated for the DOE Office of Science by Argonne National Laboratory under Contract No. DE-AC02-06CH11357. Extraordinary facility operations were supported in part by the DOE Office of Science through the National Virtual Biotechnology Laboratory, a consortium of DOE national laboratories focused on the response to COVID-19, with funding provided by the Coronavirus CARES Act. Use of the Stanford Synchrotron Radiation Lightsource, SLAC National Accelerator Laboratory, is supported by the U.S. Department of Energy, Office of Science, Office of Basic Energy Sciences under Contract No. DE-AC02-76SF00515. The SSRL Structural Molecular Biology Program is supported by the DOE Office of Biological and Environmental Research, and by the National Institutes of Health, National Institute of General Medical Sciences (P30GM133894).

## Author Contributions

P.Z., M.Y., Y.Z., O.L., J.G.J.,I.A.W., and R.A. conceived and designed the study. N.B. and T.F.R. recruited donors and collected and processed plasma samples. P.Z., W.-t.H., and G.S. perfomed single B cell sorting. P.Z., Y.Z., and T.C. carried out neutralization assays. P.Z., Y.Z., G.A., and P.Y. perfomed BLI and ELISA. P.Z. and X.L. did CELISA binding. P.Z., S.C., and F.A. generated recombinant soluble spike proteins. P.Z. and O.L. perfomed HEp2 polyreactive assay abd PSR ELISA. M.Y., H.L., and I.A.W. determined the crystal structure of the antibody-antigen complex. P.Z., O.L., F.Z., and J.G.J. designed the synthetic antibody library and performed the yeast library display and FACS sorting. P.Z., M.Y., Y.Z., O.L., D.R.B., J.G.J.,I.A.W., and R.A. designed the experiments and analyzed the data. P.Z., M.Y., Y.Z., J.G.J.,I.A.W., and R.A. wrote the paper. All authors reviewed and edited the paper.

## Competing Interests

All authors have declared that no competing interests exist.

## Data Availability

All data associated with this study are present in the paper or the supplementary materials. The structures reported in this study are available under the accession codes PDB ID XXXX, XXXX and XXXX.

---

**Fig S1.**
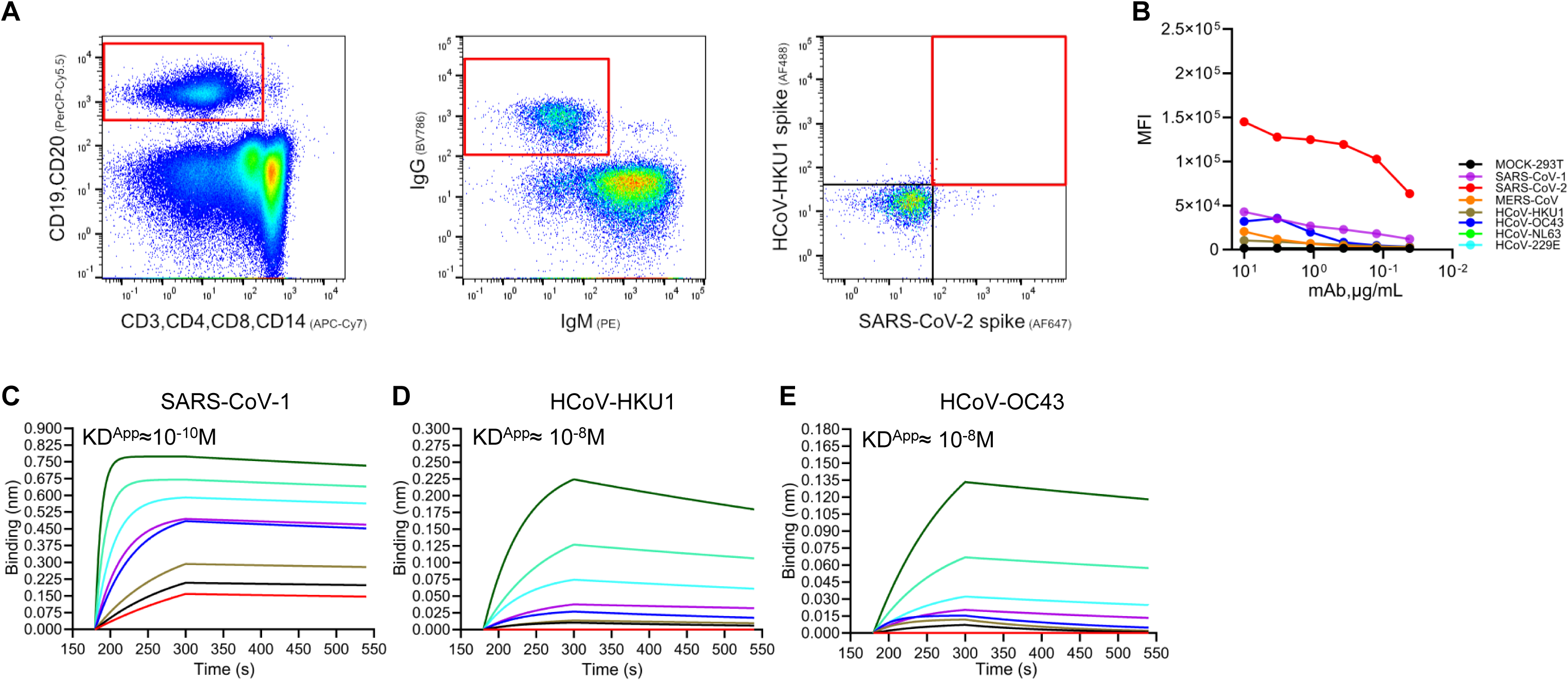
B cell sorting strategy and binding ability of parental CC65.1. (**A**) Gating strategy of single cross-reactive lgG^+^ B cell sorting from CC65 donor. The CD19^+^CD20^+^CD3^−^CD4^−^CD8^−^CD14^−^IgM^−^IgG^+^HCoV-HKU1^+^SARS-CoV2^+^ cells were sorted. (**B**) Binding of CC65.1 to the cell surface-expressed spikes from different alphacoronaviruses and betacoronaviruses was test by CELISA, mean fluorescence intensity (MFI) was shown. (**C-E**) BLI binding curves of CC65.1 with SARS-CoV-1 (C), HCoV-HKU1 (D), and HCoV-OC43 (E) recombinant soluble spike proteins by BLI. The KD^APP^ is indicated.

**Fig S2.**
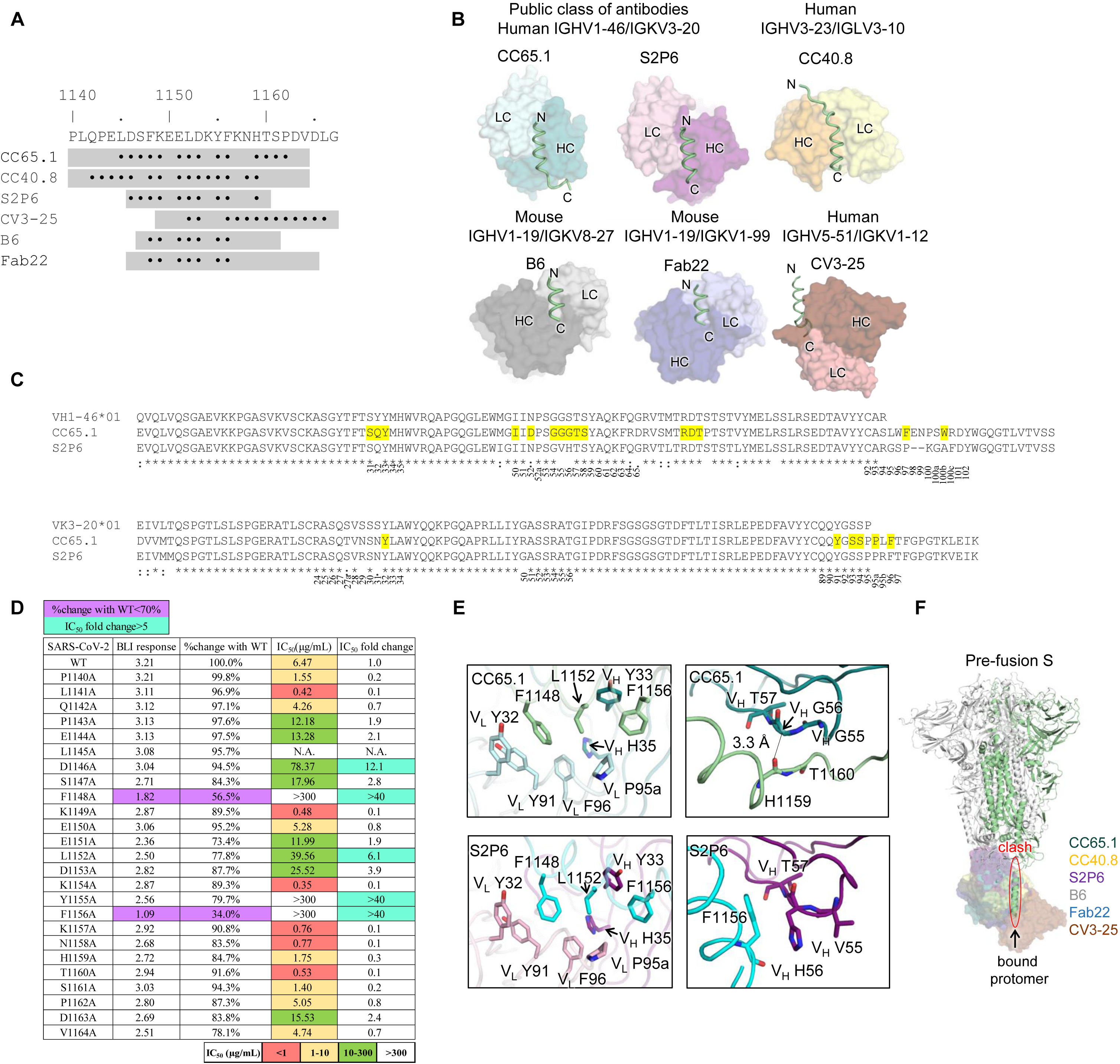
Comparison of S2 stem helix targeting antibodies. (**A**) Epitope residues (buried surface area, BSA > 0 Å^2^) of each antibody are indicated by dots under the sequence. Peptides used for each crystallization study are indicated as gray boxes. (**B**) Structures of CC65.1, CC40.8, S2P6, B6, CV3-25, and Fab22 in complex with SARS-CoV-1/2 S2 stem peptides. The SARS-CoV-1/2 S2 stem helices are represented by green tubes. N- and C-terminus of the bound antigens are indicated in the figure. (**C**) Sequence alignment between CC65.1, S2P6, and their putative germline sequences IGHV1-46*01 and IGKV3-20*01. Paratope residues shown in Fig S2E are highlighted in yellow. Residues conserved in all aligned sequences are labelled by an asterisk (*), whereas a colon (:) and a period (.) indicate strongly similar and weakly similar sequences, respectively as calculated by Clustal Omega [46]. Kabat numbering is shown under the sequence alignment. (**D**) Epitope mapping of CC65.1 by alanine scanning mutagenesis of SARS-CoV-1/2 stem helix peptides using BLI and neutralization assays. The IC_50_ fold change (n-fold) was calculated by dividing the mutant value by the WT value. For IC50, n-fold >5 is indicated as cyan. For binding response values where the % change in binding (compared to WT peptide) is <70%, are indicated in purple. N.A., not available. (**E**) Structural comparisons of conserved (left) and non-conserved (right) interactions with the SARS-CoV-1/2 stem helix between the two IGHV1-46*01/IGKV3-20*01 antibodies CC65.1 and S2P6. (**F**) SARS-CoV-2 spike pre-fusion structure (PDB 6XR8) is superimposed with the structures of CC65.1 (teal) CC40.8 (yellow), S2P6 (purple), B6 (grey), CV3-25 (brown), and Fab22 (blue) in complex with SARS-CoV-2 S2 peptides. All antibodies would clash with the other protomers of the spike protein in pre-fusion state. Structures of CC40.8, S2P6, B6, CV3-25 and Fab22 are from PDBs 7SJS, 7RNJ, 7M53, 7NAB, and 7S3N, respectively.

**Fig S3.**
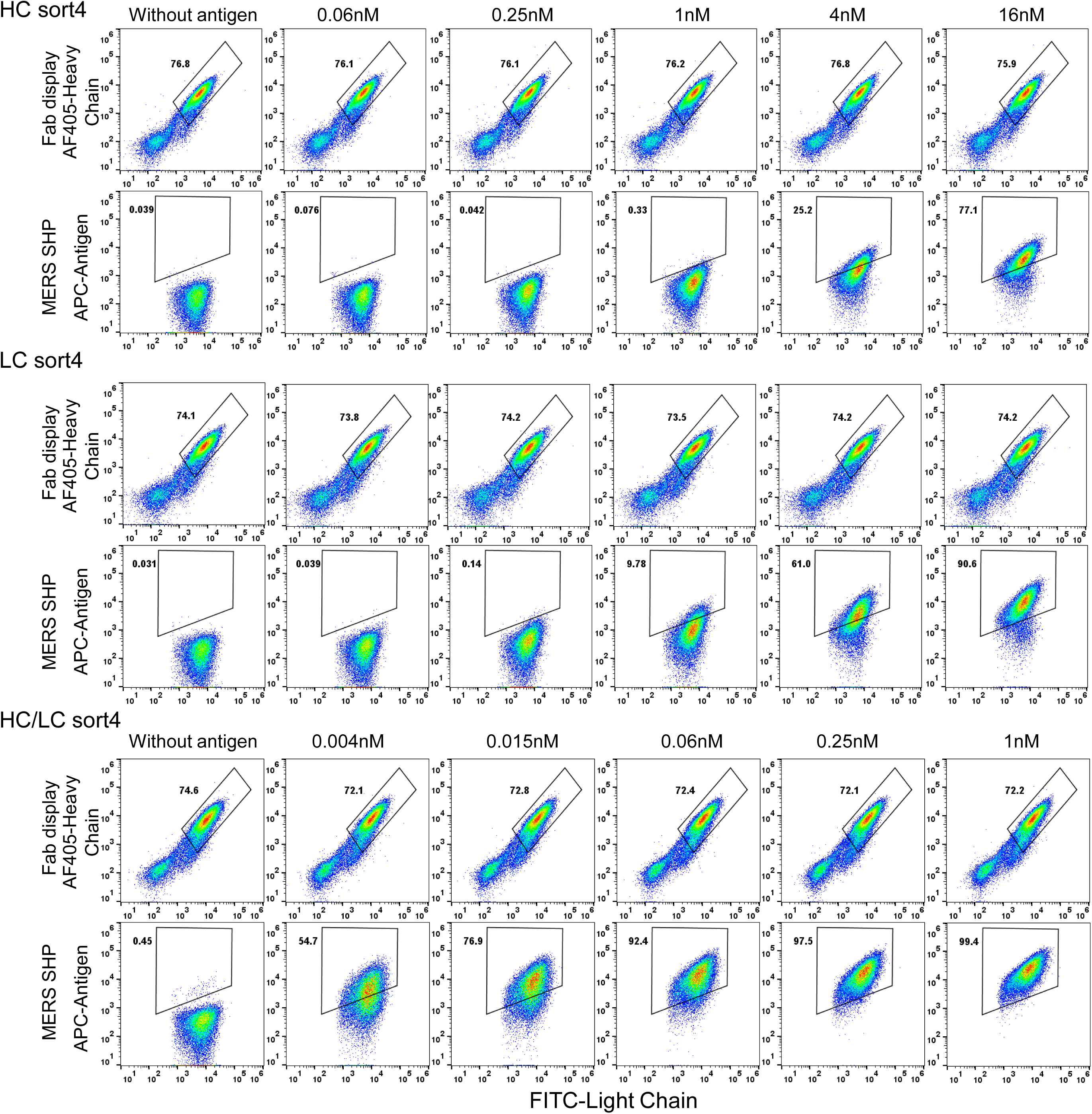
Representative FACS plots of CC65.1 HC, LC and HC/LC libraries in 4^th^ sorting. Surface Fab display frequency was determined by staining with AF405-anti-V5 antibody (HC) and FITC-anti-c-Myc (LC). Cells were also labeled with different unsaturated concentrations of biotinylated MERS-CoV stem helix peptide, or without antigen. Labeled cells were further stained with APC conjugated streptavidin. FACS analysis was performed by FlowJo. SHP, stem helix peptide.

**Fig S4.**
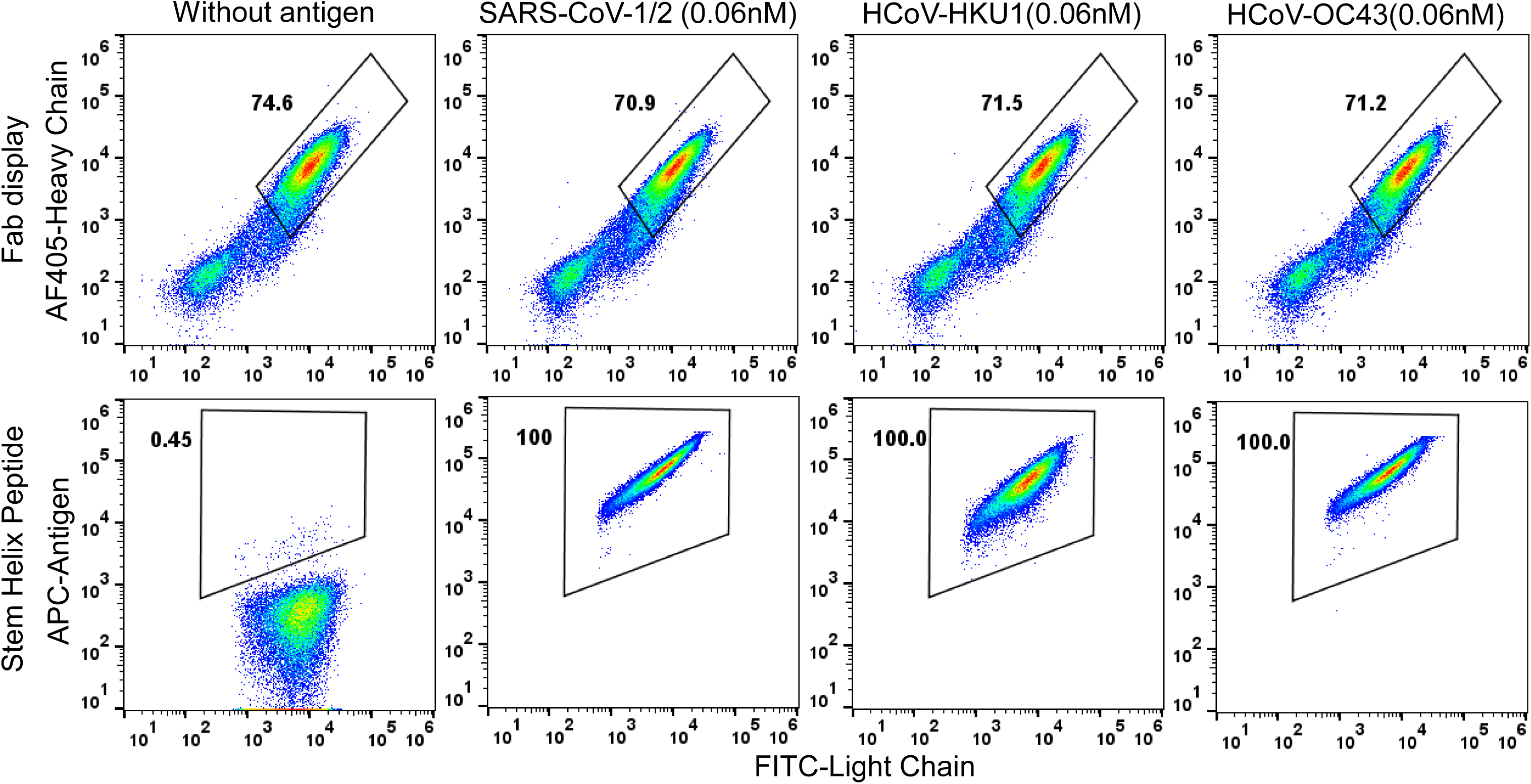
Binding of yeast cells in the 4^th^ sort of HC/LC library to stem helix peptides of SARS-CoV-1/2, HCoV-HKU1, and HCoV-OC43. AF405-anti-V5 antibody (HC) and FITC-anti-c-Myc (LC) were used in combination to determine the cell surface Fab display frequency. Cells were also labeled with different biotinylated betacoronavirus stem helix peptides at unsaturated concentration (0.06 nM), or without antigen. Labeled cells were further stained with APC conjugated streptavidin. FACS analysis was performed by FlowJo.

**Fig S5.**
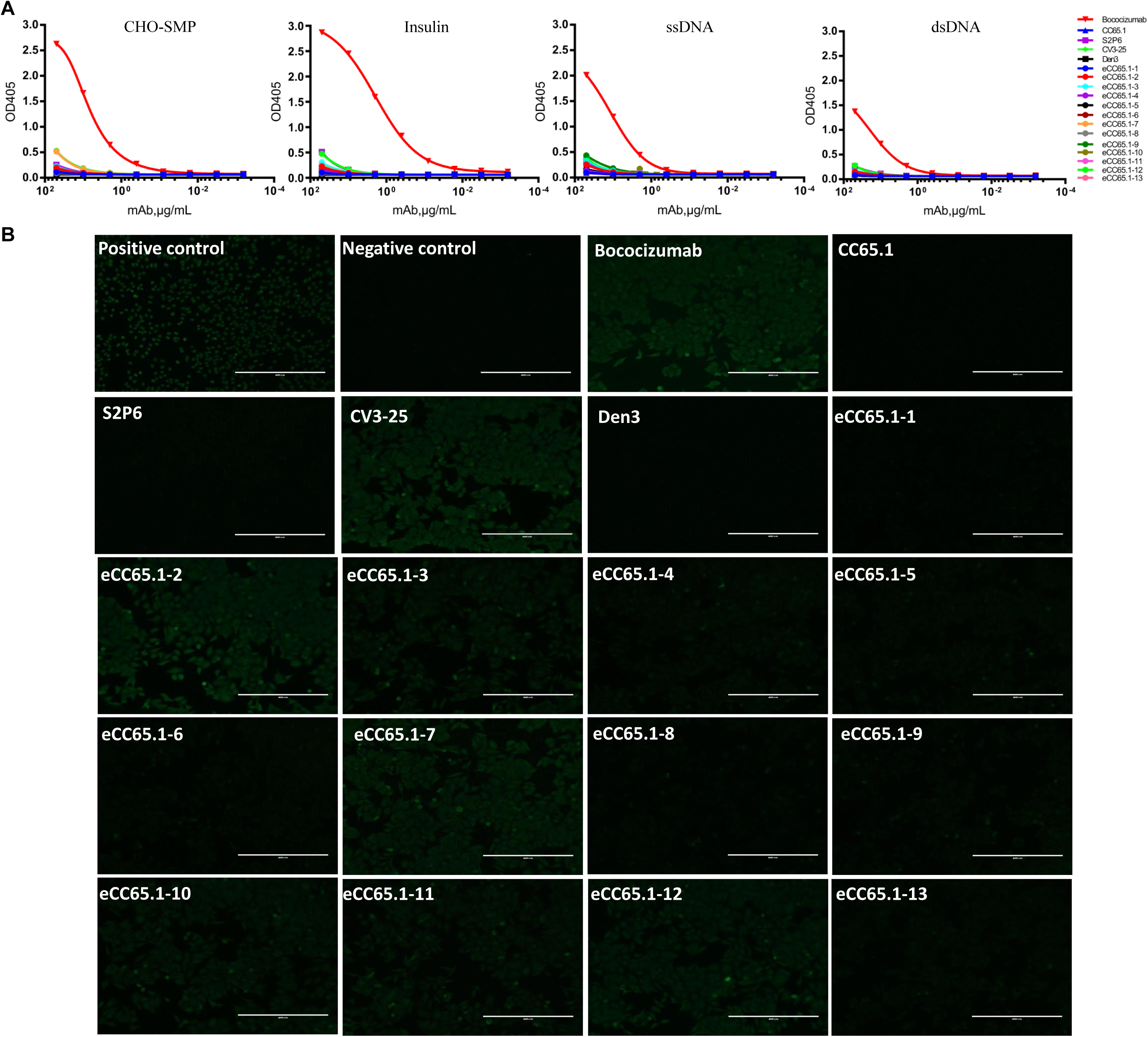
Evaluation of selected eCC65.1 mAbs for polyreactivity. eCC65.1 mAbs were tested (**A**) by ELISA for binding against polyspecific reagents (PSR) including Chinese hamster ovary cells solubilized membrane protein (CHO-SMP), insulin, single-strand DNA (ssDNA) and double-strand DNA (dsDNA) and (**B**) by binding to immobilized HEp2 epithelial cells. Bococizumab which used as a positive control is a humanized mAb targeting the LDL receptor-binding domain of PCSK9 and it has been studied in phase I–III clinical studies [47]. Den3 was used as a negative control. Positive and negative controls for the HEp2 kit assay were provided by the manufacturer.

**Fig S6.**
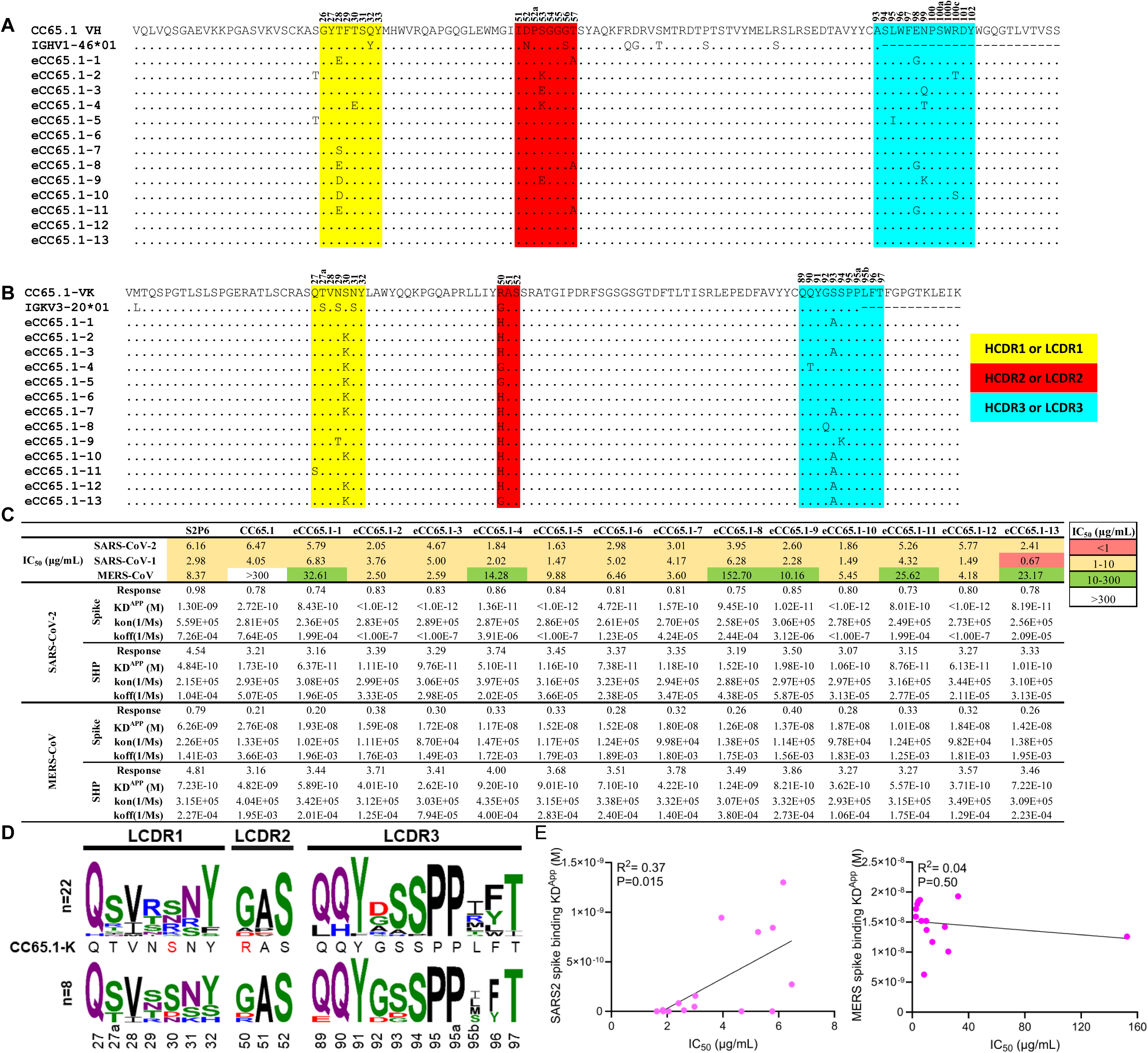
**Sequence analysis, binding affinity, and neutralizing potency of selected eCC65.1 mAbs**. Variable regions of heavy chains (**A**) and light chains (**B**) of CC65.1, selected eCC65.1 mAbs and their putative germline sequences IGHV1-46*01 and IGKV3-20*01 were aligned, CDRs are highlighted in different colors, yellow for HCDR1 and LCDR1, red for HCDR2 and LCDR2 and cyan for HCDR3 and LCDR3. Kabat numbering is shown above the sequence alignment (**C**) Summary table of binding affinity and neutralizing potency of parental CC65.1 and eCC65.1 mAbs. IC_50_ of CC65.1, eCC65.1 mAbs, and S2P6 are shown in the upper table. Binding affinity of these antibodies to recombinant soluble spikes and stem helix peptides of SARS-CoV-2 and MERS-CoV were tested by BLI, values of response, KD^APP^, kon and koff were analyzed by fitting to 1:1 model. (**D**) Identification of signature residues on LCDRs between natural occurring KV3-20- derived MERS-CoV-neutralizing (top, n=22) and MERS-CoV-non-neutralizing (bottom, n=8) human stem helix mAbs, sequences were aligned and sequence logo of LCDRs is shown. Sequences of CC65.1 LCDRs are shown, the signature substitutions enriched among the eCC65.1 antibodies are highlighted in red. Kabat numbering is shown under the sequence alignment. KV3-20-derived MERS-CoV-neutralizing human stem helix bnAbs include S2P6 [18], CC9.104, CC9.106, CC9.111, CC9.113, CC9.130, CC9.131, CC24.107, CC25.101, CC25.104, CC68.104, CC68.109, CC92.133, CC92.147, CC99.103, CC99.104, CC99.105[13], COV89-22, COV30-14, COV72-37, COV44-26 and COV44-74 [19]. KV3-20-derived MERS-CoV-non-neutralizing human stem helix mAbs contain CC65.1, CC24.105, CC25.108, CC25.112, CC67.105, CC67.130, CC95.102 [13] and COV93-03 [19]. Sequences are indicated by colors representing their different biochemical properties: green for polar, blue for basic, red for acidic, black for hydrophobic and purple for N or Q residues. (**E**) Analysis of possible correlations of S2P6, CC65.1 and selected eCC65.1 mAbs’ binding affinity (KD^APP^) to SARS-CoV-2 and MERS-CoV recombinant soluble spikes with neutralization potency (IC_50_) against their corresponding viruses. Correlations were determined by nonparametric Spearman correlation two-tailed test with 95% confidence interval. The Spearman correlation coefficient (r squared) and p-values are indicated.

**Fig S7.**
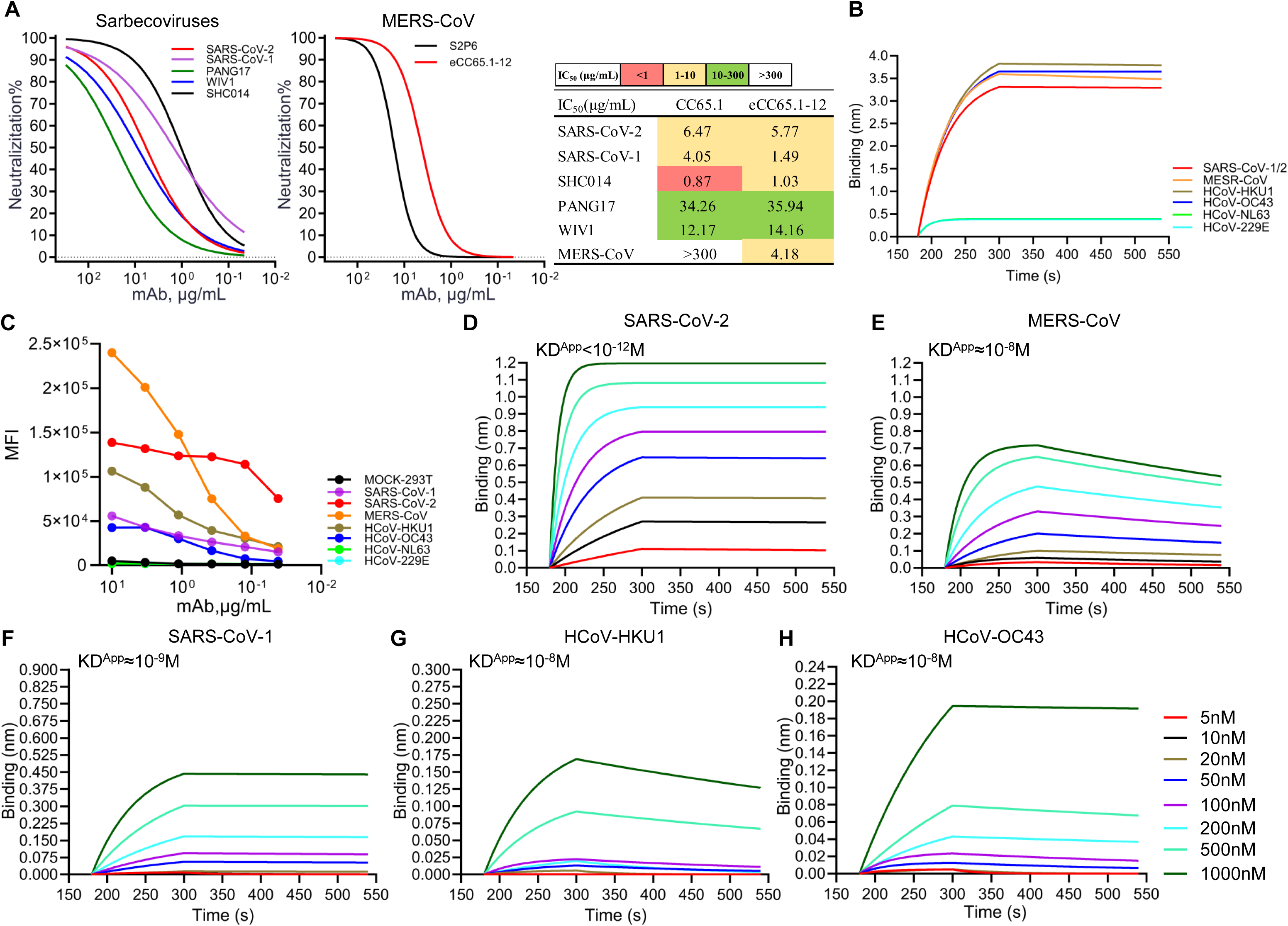
Neutralizing potency and binding ability of eCC65.1-12 to betacoronaviruses. (**A**) Neutralization potency of eCC65.1-12 to sarbecoviruses (left) and MERS-CoV (right), with S2P6 as a positive control. Table showing the IC_50_ of CC65.1 and eCC65.1-12 to sarbecoviruses and MERS-CoV. (**B**) BLI binding of eCC65.1-12 to 25-mer stem helix peptides derived from different coronavirus spikes. (**C**) Binding of eCC65.1-12 mAb with betacoronavirus and alphacoronavirus spikes expressed on the HEK293T cell surface. MFI, Mean Fluorescence Intensity. (**D-H**) BLI binding curves of eCC65.1-12 with all the five human betacoronavirus recombinant soluble spike proteins. eCC65.1-12 was captured on a Protein A biosensor, followed by exposure to varying concentrations of spike protein. The KD^App^ was determined using a 1:1 binding kinetics model with ForteBio Data Analysis software.

**Table S1.** Summary table of mean fluorescence intensity (MFI) and binding affinity of parental CC65.1 and eCC65.1-12 mAbs.

|  |  | MFI |  |  |  |  |  | Spike |  |  | Stem helix peptides |  |  |
| --- | --- | --- | --- | --- | --- | --- | --- | --- | --- | --- | --- | --- | --- |
|  |  | 10.00 | 3.33 | 1.11 | 0.37 | 0.12 | 0.04 | KD<sup>APP</sup> (M) | kon (1/Ms) | koff (1/Ms) | KD<sup>APP</sup> (M) | kon (1/Ms) | koff (1/Ms) |
| CC65.1 | MOCK-293T | 1474 | 1579 | 1430 | 1534 | 1493 | 1376 | N.D. |  |  | N.D. |  |  |
|  | SARS-CoV-1 | 42857 | 34906 | 26742 | 22798 | 18074 | 11906 | 8.20E-10 | 4.32E+05 | 3.54E-04 | 1.73E-10 | 2.93E+05 | 5.07E-05 |
|  | SARS-CoV-2 | 144905 | 127685 | 124789 | 119288 | 102680 | 63400 | 2.72E-10 | 2.81E+05 | 7.64E-05 |  |  |  |
|  | MERS-CoV | 20548 | 11837 | 6773 | 3893 | 2549 | 2003 | 2.76E-08 | 1.33E+05 | 3.66E-03 | 4.82E-09 | 4.04E+05 | 1.95E-03 |
|  | HCoV-HKU1 | 10345 | 8964 | 7060 | 5233 | 3839 | 2979 | 2.12E-08 | 1.16E+05 | 2.46E-03 | 8.67E-10 | 2.45E+05 | 2.13E-04 |
|  | HCoV-OC43 | 31898 | 35505 | 19859 | 8288 | 4749 | 3280 | 3.08E-08 | 5.77E+04 | 1.77E-03 | 1.24E-10 | 2.93E+05 | 3.62E-05 |
|  | HCoV-NL63 | 1640 | 1623 | 1623 | 1651 | 1621 | 1613 | N.D. |  |  | N.D. |  |  |
|  | HCoV-229E | 1642 | 1620 | 1603 | 1634 | 1624 | 1621 | N.D. |  |  | N.D. |  |  |
| CC65.1-12 | MOCK-293T | 4793 | 3251 | 1759 | 1533 | 1430 | 1407 | N.D. |  |  | N.D. |  |  |
|  | SARS-CoV-1 | 55745 | 42960 | 33492 | 26370 | 20731 | 15207 | 2.03E-09 | 5.26E+04 | 1.07E-04 | 6.13E-11 | 3.44E+05 | 2.11E-05 |
|  | SARS-CoV-2 | 138682 | 131730 | 123676 | 122520 | 114021 | 75275 | <1.00E-12 | 2.73E+05 | <1.00E-7 |  |  |  |
|  | MERS-CoV | 240163 | 200902 | 147890 | 75072 | 32996 | 18774 | 1.84E-08 | 9.82E+04 | 1.81E-03 | 3.71E-10 | 3.49E+05 | 1.29E-04 |
|  | HCoV-HKU1 | 106435 | 87983 | 56750 | 39216 | 30306 | 21206 | 5.39E-08 | 7.34E+04 | 3.95E-03 | 1.29E-10 | 3.01E+05 | 3.90E-05 |
|  | HCoV-OC43 | 42606 | 42796 | 30132 | 16550 | 7461 | 4519 | 2.92E-08 | 6.94E+04 | 2.03E-03 | 1.63E-11 | 3.26E+05 | 5.31E-06 |
|  | HCoV-NL63 | 2009 | 1877 | 1715 | 1681 | 1623 | 1613 | N.D. |  |  | N.D. |  |  |
|  | HCoV-229E | 2096 | 1834 | 1732 | 1680 | 1615 | 1627 | N.D. |  |  | N.D. |  |  |

Mean Fluorescence Intensity (MFI) of CC65.1 and eCC65.1-12 to betacoronavirus and alphacoronavirus spikes expressed on HEK293T cell surface at different mAb concentrations. Mock-transfected HEK293T cells were used as a negative control, MFI was calculated by FlowJo. Binding affinity of these antibodies to recombinant soluble spikes and stem helix peptides of betacoronavirus were tested by BLI, KD<sup>APP</sup>, kon and koff were analyzed by fitting to 1:1 model. N.D., not detected.

**Table S2.** X-ray data collection and refinement statistics

| **Data collection** |  |  |  |
| --- | --- | --- | --- |
|  | CC65.1 scFv + SARS-CoV-2 S2 stem peptide | CC65.1 Fab + MERS-CoV S2 stem peptide | eCC65.1-12 Fab + MERS-CoV S2 stem peptide |
| Beamline | SSRL 12-1 | ALS 5.0.1 | APS23-ID-B |
| Wavelength (Å) | 0.97946 | 0.97741 | 0.97930 |
| Space group | C 2 2 2<sub>1</sub> | C 2 | C 2 |
| Unit cell parameters |  |  |  |
| a, b, c (Å) | 83.5, 98.5, 144.0 | 95.4, 58.6, 96.7 | 91.0, 60.0, 81.8 |
| α, β, γ (°) | 90, 90, 90 | 90, 93.5, 90 | 90, 98.0, 90 |
| Resolution (Å)<sup>a</sup> | 50.0–2.05 (2.09–2.05) | 50.0–2.70 (2.83–2.70) | 50.0–2.51 (2.55–2.51) |
| Unique reflections<sup>a</sup> | 34,344 (2,176) | 14,853 (1,482) | 13,990 (913) |
| Redundancy<sup>a</sup> | 7.0 (2.8) | 6.5 (6.7) | 3.9 (3.4) |
| Completeness (%)<sup>a</sup> | 91.4 (55.5) | 100 (100) | 99.5 (96.3) |
| <I/σI><sup>a</sup> | 21.0 (4.4) | 5.0 (1.4) | 8.7 (1.1) |
| R<sub>sym</sub><sup>b</sup> (%)<sup>a</sup> | 13.4 (24.8) | 29.3 (>100) | 14.2 (>100) |
| R<sub>pim</sub><sup>b</sup> (%)<sup>a</sup> | 5.1 (16.0) | 18.8 (78.7) | 8.0 (59.3) |
| CC<sub>1/2</sub><sup>c</sup> (%)<sup>a</sup> | 98.4 (92.6) | 97.5 (63.6) | 97.5 (50.0) |
| **Refinement statistics** |  |  |  |
| Resolution (Å) | 40.1–2.05 | 48.3–2.70 | 44.0–2.51 |
| Reflections (work) | 34,341 | 14,833 | 13,988 |
| Reflections (test) | 2,177 | 1,479 | 913 |
| R<sub>cryst</sub><sup>d</sup> / R<sub>free</sub><sup>e</sup> (%) | 19.2/22.9 | 25.0/30.6 | 21.8/26.4 |
| No. of atoms | 4,375 | 3,562 | 3,600 |
| Antibodies | 3,562 | 3,314 | 3,353 |
| Peptides | 322 | 141 | 113 |
| Other (SO4) | 0 | 20 | 25 |
| Solvent | 491 | 87 | 109 |
| Average B-value (Å<sup>2</sup>) | 26 | 29 | 43 |
| Antibodies | 24 | 29 | 42 |
| Peptide | 38 | 38 | 59 |
| Other (SO4) | N/A | 36 | 73 |
| Solvent | 31 | 26 | 37 |
| Wilson B-value (Å<sup>2</sup>) | 21 | 33 | 43 |
| **RMSD from ideal geometry** |  |  |  |
| Bond length (Å) | 0.002 | 0.003 | 0.02 |
| Bond angle (°) | 0.55 | 0.71 | 0.58 |
| **Ramachandran statistics<sup>f</sup>** |  |  |  |
| Favored | 97.5 | 93.7 | 96.0 |
| Outliers | 0 | 0 | 0 |
| **PDB code** | pending | pending | pending |

<sup>a</sup> Numbers in parentheses refer to the highest resolution shell.
<sup>b</sup> $R_{\text{sym}} = \sum_{hkl} \sum_i |I_{hkl,i} - \langle I_{hkl} \rangle| / \sum_{hkl} \sum_i I_{hkl,i}$ and $R_{\text{pim}} = \sum_{hkl} (1/(n-1))^{1/2} \sum_i |I_{hkl,i} - \langle I_{hkl} \rangle| / \sum_{hkl} \sum_i I_{hkl,i}$ , where $I_{hkl,i}$ is the scaled intensity of the $i^{\text{th}}$ measurement of reflection h, k, l, $\langle I_{hkl} \rangle$ is the average intensity for that reflection, and $n$ is the redundancy.
<sup>c</sup> CC<sub>1/2</sub> = Pearson correlation coefficient between two random half datasets.
<sup>d</sup> $R_{\text{cryst}} = \sum_{hkl} |F_o - F_c| / \sum_{hkl} |F_o| \times 100$ , where $F_o$ and $F_c$ are the observed and calculated structure factors, respectively.
<sup>e</sup> $R_{\text{free}}$ was calculated as for $R_{\text{cryst}}$ , but on a test set comprising 5% of the data excluded from refinement.
<sup>f</sup> From MolProbity [48].

## Notes

### Competing Interest Statement

The authors have declared no competing interest.

## References

1. Cui J, Li F, Shi ZL. Origin and evolution of pathogenic coronaviruses. Nat Rev Microbiol. 2019;17(3):181–92. doi: 10.1038/s41579-018-0118-9. PubMed PMID: 30531947; PubMed Central PMCID: PMCPMC7097006.

2. Wang Q, Guo Y, Mellis IA, Wu M, Mohri H, Gherasim C, et al. Antibody evasiveness of SARS-CoV-2subvariants KP.3.1.1 and XEC. Cell Rep. 2025;44(4):115543. Epub 20250408. doi: 10.1016/j.celrep.2025.115543. PubMed PMID: 40202847; PubMed Central PMCID: PMCPMC12014523.

3. Wang Q, Iketani S, Li Z, Liu L, Guo Y, Huang Y, et al. Alarming antibody evasion properties of rising SARS-CoV-2 BQ and XBB subvariants. Cell. 2023;186(2):279–86.e8. Epub 20221214. doi: 10.1016/j.cell.2022.12.018. PubMed PMID: 36580913; PubMed Central PMCID: PMCPMC9747694.

4. Wec AZ, Wrapp D, Herbert AS, Maurer DP, Haslwanter D, Sakharkar M, et al. Broad neutralization of SARS-related viruses by human monoclonal antibodies. Science. 2020;369(6504):731-6. Epub 20200615. doi: 10.1126/science.abc7424. PubMed PMID: 32540900; PubMed Central PMCID: PMCPMC7299279.

5. Liu L, Iketani S, Guo Y, Chan JF, Wang M, Liu L, et al. Striking antibody evasion manifested by the Omicron variant of SARS-CoV-2. Nature. 2022;602(7898):676-81. Epub 20211223. doi: 10.1038/s41586-021-04388-0. PubMed PMID: 35016198.

6. Peiris M, Perlman S. Unresolved questions in the zoonotic transmission of MERS. Curr Opin Virol. 2022;52:258–64. Epub 20220106. doi: 10.1016/j.coviro.2021.12.013. PubMed PMID: 34999369; PubMed Central PMCID: PMCPMC8734234.

7. Anthony SJ, Gilardi K, Menachery VD, Goldstein T, Ssebide B, Mbabazi R, et al. Further evidence for bats as the evolutionary source of Middle East Respiratory Syndrome coronavirus.

8. mBio. 2017;8(2):e00373-17. Epub 20170404. doi: 10.1128/mBio.00373-17. PubMed PMID: 28377531; PubMed Central PMCID: PMCPMC5380844.

8. de Wit E, van Doremalen N, Falzarano D, Munster VJ. SARS and MERS: recent insights into emerging coronaviruses. Nat Rev Microbiol. 2016;14(8):523–34. Epub 20160627. doi: 10.1038/nrmicro.2016.81. PubMed PMID: 27344959; PubMed Central PMCID: PMCPMC7097822.

9. Zaki AM, van Boheemen S, Bestebroer TM, Osterhaus AD, Fouchier RA. Isolation of a novel coronavirus from a man with pneumonia in Saudi Arabia. N Engl J Med. 2012;367(19):1814–20. Epub 20121017. doi: 10.1056/NEJMoa1211721. PubMed PMID: 23075143.

10. Greaney AJ, Starr TN, Gilchuk P, Zost SJ, Binshtein E, Loes AN, et al. Complete mapping of mutations to the SARS-CoV-2 spike receptor-binding domain that escape antibody recognition. Cell Host Microbe. 2021;29(1):44–57.e9. Epub 20201119. doi: 10.1016/j.chom.2020.11.007. PubMed PMID: 33259788; PubMed Central PMCID: PMCPMC7676316.

11. Lu R, Zhao X, Li J, Niu P, Yang B, Wu H, et al. Genomic characterisation and epidemiology of 2019 novel coronavirus: implications for virus origins and receptor binding. Lancet. 2020;395(10224):565-74. Epub 20200130. doi: 10.1016/s0140-6736(20)30251-8. PubMed PMID: 32007145; PubMed Central PMCID: PMCPMC7159086.

12. Menachery VD, Yount BL, Jr., Sims AC, Debbink K, Agnihothram SS, Gralinski LE, et al. SARS-like WIV1-CoV poised for human emergence. Proc Natl Acad Sci U S A. 2016;113(11):3048–53. Epub 20160314. doi: 10.1073/pnas.1517719113. PubMed PMID: 26976607; PubMed Central PMCID: PMCPMC4801244.

13. Zhou P, Song G, Liu H, Yuan M, He WT, Beutler N, et al. Broadly neutralizing anti-S2 antibodies protect against all three human betacoronaviruses that cause deadly disease. Immunity. 2023;56(3):669–86.e7. Epub 20230216. doi: 10.1016/j.immuni.2023.02.005. PubMed PMID: 36889306; PubMed Central PMCID: PMCPMC9933850.

14. Song G, He WT, Callaghan S, Anzanello F, Huang D, Ricketts J, et al. Cross-reactive serum and memory B-cell responses to spike protein in SARS-CoV-2 and endemic coronavirus infection. Nat Commun. 2021;12(1):2938. Epub 20210519. doi: 10.1038/s41467-021-23074-3. PubMed PMID: 34011939; PubMed Central PMCID: PMCPMC8134462.

15. Liu H, Wilson IA. Protective neutralizing epitopes in SARS-CoV-2. Immunol Rev. 2022;310(1):76–92. Epub 20220522. doi: 10.1111/imr.13084. PubMed PMID: 35599305; PubMed Central PMCID: PMCPMC9348472.

16. Parren PW, Burton DR. The antiviral activity of antibodies in vitro and in vivo. Adv Immunol. 2001;77:195–262. doi: 10.1016/s0065-2776(01)77018-6. PubMed PMID: 11293117; PubMed Central PMCID: PMCPMC7131048.

17. Rappazzo CG, Tse LV, Kaku CI, Wrapp D, Sakharkar M, Huang D, et al. Broad and potent activity against SARS-like viruses by an engineered human monoclonal antibody. Science. 2021;371(6531):823-9. Epub 20210125. doi: 10.1126/science.abf4830. PubMed PMID: 33495307; PubMed Central PMCID: PMCPMC7963221.

18. Pinto D, Sauer MM, Czudnochowski N, Low JS, Tortorici MA, Housley MP, et al. Broad betacoronavirus neutralization by a stem helix-specific human antibody. Science. 2021;373(6559):1109-16. Epub 20210806. doi: 10.1126/science.abj3321. PubMed PMID: 34344823; PubMed Central PMCID: PMCPMC9268357.

19. Dacon C, Peng L, Lin TH, Tucker C, Lee CD, Cong Y, et al. Rare, convergent antibodies targeting the stem helix broadly neutralize diverse betacoronaviruses. Cell Host Microbe. 2023;31(1):97–111.e12. Epub 20221107. doi: 10.1016/j.chom.2022.10.010. PubMed PMID: 36347257; PubMed Central PMCID: PMCPMC9639329.

20. Zhou P, Yuan M, Song G, Beutler N, Shaabani N, Huang D, et al. A human antibody reveals a conserved site on beta-coronavirus spike proteins and confers protection against SARS-CoV-2 infection. Sci Transl Med. 2022;14(637):eabi9215. Epub 20220323. doi: 10.1126/scitranslmed.abi9215. PubMed PMID: 35133175; PubMed Central PMCID: PMCPMC8939767.

21. Jennewein MF, MacCamy AJ, Akins NR, Feng J, Homad LJ, Hurlburt NK, et al. Isolation and characterization of cross-neutralizing coronavirus antibodies from COVID-19+ subjects. Cell Rep. 2021;36(2):109353. Epub 20210622. doi: 10.1016/j.celrep.2021.109353. PubMed PMID: 34237283; PubMed Central PMCID: PMCPMC8216847.

22. Li W, Chen Y, Prévost J, Ullah I, Lu M, Gong SY, et al. Structural basis and mode of action for two broadly neutralizing antibodies against SARS-CoV-2 emerging variants of concern. Cell Rep. 2022;38(2):110210. Epub 20211215. doi: 10.1016/j.celrep.2021.110210. PubMed PMID: 34971573; PubMed Central PMCID: PMCPMC8673750.

23. Sauer MM, Tortorici MA, Park YJ, Walls AC, Homad L, Acton OJ, et al. Structural basis for broad coronavirus neutralization. Nat Struct Mol Biol. 2021;28(6):478–86. Epub 20210512. doi: 10.1038/s41594-021-00596-4. PubMed PMID: 33981021.

24. Hsieh CL, Werner AP, Leist SR, Stevens LJ, Falconer E, Goldsmith JA, et al. Stabilized coronavirus spike stem elicits a broadly protective antibody. Cell Rep. 2021;37(5):109929. Epub 20211016. doi: 10.1016/j.celrep.2021.109929. PubMed PMID: 34710354; PubMed Central PMCID: PMCPMC8519809.

25. Zhou P, Wang H, Fang M, Li Y, Wang H, Shi S, et al. Broadly resistant HIV-1 against CD4- binding site neutralizing antibodies. PLoS Pathog. 2019;15(6):e1007819. Epub 20190613. doi: 10.1371/journal.ppat.1007819. PubMed PMID: 31194843; PubMed Central PMCID: PMCPMC6592578.

26. Zhao F, Yuan M, Keating C, Shaabani N, Limbo O, Joyce C, et al. Broadening a SARS- CoV-1-neutralizing antibody for potent SARS-CoV-2 neutralization through directed evolution. Sci Signal. 2023;16(798):eabk3516. Epub 20230815. doi: 10.1126/scisignal.abk3516. PubMed PMID: 37582161; PubMed Central PMCID: PMCPMC11771511.

27. Julian MC, Li L, Garde S, Wilen R, Tessier PM. Efficient affinity maturation of antibody variable domains requires co-selection of compensatory mutations to maintain thermodynamic stability. Sci Rep. 2017;7:45259. Epub 20170328. doi: 10.1038/srep45259. PubMed PMID: 28349921; PubMed Central PMCID: PMCPMC5368667.

28. Zhao F, Keating C, Ozorowski G, Shaabani N, Francino-Urdaniz IM, Barman S, et al. Engineering SARS-CoV-2 neutralizing antibodies for increased potency and reduced viral escape pathways. iScience. 2022;25(9):104914. Epub 20220811. doi: 10.1016/j.isci.2022.104914. PubMed PMID: 35971553; PubMed Central PMCID: PMCPMC9367177.

29. Casadevall A, Focosi D. Lessons from the use of monoclonal antibodies to SARS-CoV-2 spike protein during the COVID-19 pandemic. Annu Rev Med. 2025;76(1):1–12. Epub 20250116. doi: 10.1146/annurev-med-061323-073837. PubMed PMID: 39630849.

30. Cohen AA, Gnanapragasam PNP, Lee YE, Hoffman PR, Ou S, Kakutani LM, et al. Mosaic nanoparticles elicit cross-reactive immune responses to zoonotic coronaviruses in mice. Science. 2021;371(6530):735-41. Epub 20210112. doi: 10.1126/science.abf6840. PubMed PMID: 33436524; PubMed Central PMCID: PMCPMC7928838.

31. Hurlburt NK, Homad LJ, Sinha I, Jennewein MF, MacCamy AJ, Wan YH, et al. Structural definition of a pan-sarbecovirus neutralizing epitope on the spike S2 subunit. Commun Biol. 2022;5(1):342. Epub 20220411. doi: 10.1038/s42003-022-03262-7. PubMed PMID: 35411021; PubMed Central PMCID: PMCPMC9001700.

32. Li CJ, Chang SC. SARS-CoV-2 spike S2-specific neutralizing antibodies. Emerg Microbes Infect. 2023;12(2):2220582. doi: 10.1080/22221751.2023.2220582. PubMed PMID: 37254830; PubMed Central PMCID: PMCPMC10274517.

33. Edwards CT, Karunakaran KA, Garcia E, Beutler N, Gagne M, Golden N, et al. Passive infusion of an S2-Stem broadly neutralizing antibody protects against SARS-CoV-2 infection and lower airway inflammation in rhesus macaques. PLoS Pathog. 2025;21(1):e1012456. Epub 20250123. doi: 10.1371/journal.ppat.1012456. PubMed PMID: 39847599; PubMed Central PMCID: PMCPMC11793774.

34. Harvey WT, Carabelli AM, Jackson B, Gupta RK, Thomson EC, Harrison EM, et al. SARS- CoV-2 variants, spike mutations and immune escape. Nat Rev Microbiol. 2021;19(7):409–24. Epub 20210601. doi: 10.1038/s41579-021-00573-0. PubMed PMID: 34075212; PubMed Central PMCID: PMCPMC8167834.

35. Letko M, Marzi A, Munster V. Functional assessment of cell entry and receptor usage for SARS-CoV-2 and other lineage B betacoronaviruses. Nat Microbiol. 2020;5(4):562–9. Epub 20200224. doi: 10.1038/s41564-020-0688-y. PubMed PMID: 32094589; PubMed Central PMCID: PMCPMC7095430.

36. Chen Y, Zhao X, Zhou H, Zhu H, Jiang S, Wang P. Broadly neutralizing antibodies to SARS-CoV-2 and other human coronaviruses. Nat Rev Immunol. 2023;23(3):189–99. Epub 20220927. doi: 10.1038/s41577-022-00784-3. PubMed PMID: 36168054; PubMed Central PMCID: PMCPMC9514166.

37. Baum A, Fulton BO, Wloga E, Copin R, Pascal KE, Russo V, et al. Antibody cocktail to SARS-CoV-2 spike protein prevents rapid mutational escape seen with individual antibodies. Science. 2020;369(6506):1014-8. Epub 20200615. doi: 10.1126/science.abd0831. PubMed PMID: 32540904; PubMed Central PMCID: PMCPMC7299283.

38. Henikoff S, Henikoff JG. Amino acid substitution matrices from protein blocks. Proc Natl Acad Sci U S A. 1992;89(22):10915–9. doi: 10.1073/pnas.89.22.10915. PubMed PMID: 1438297; PubMed Central PMCID: PMCPMC50453.

39. He WT, Musharrafieh R, Song G, Dueker K, Tse LV, Martinez DR, et al. Targeted isolation of diverse human protective broadly neutralizing antibodies against SARS-like viruses. Nat Immunol. 2022;23(6):960–70. Epub 20220602. doi: 10.1038/s41590-022-01222-1. PubMed PMID: 35654851; PubMed Central PMCID: PMCPMC10083051.

40. Khalek IS, Senji Laxme RR, Nguyen YTK, Khochare S, Patel RN, Woehl J, et al. Synthetic development of a broadly neutralizing antibody against snake venom long-chain α-neurotoxins. Sci Transl Med. 2024;16(735):eadk1867. Epub 20240221. doi: 10.1126/scitranslmed.adk1867. PubMed PMID: 38381847; PubMed Central PMCID: PMCPMC7617732.

41. Chao G, Lau WL, Hackel BJ, Sazinsky SL, Lippow SM, Wittrup KD. Isolating and engineering human antibodies using yeast surface display. Nat Protoc. 2006;1(2):755–68. doi: 10.1038/nprot.2006.94. PubMed PMID: 17406305.

42. Otwinowski Z, Minor W. Processing of X-ray diffraction data collected in oscillation mode. Methods Enzymol. 1997;276:307–26. doi: 10.1016/s0076-6879(97)76066-x. PubMed PMID: 27754618.

43. Krissinel E, Henrick K. Inference of macromolecular assemblies from crystalline state. J Mol Biol. 2007;372(3):774–97. Epub 20070513. doi: 10.1016/j.jmb.2007.05.022. PubMed PMID: 17681537.

44. Adams PD, Afonine PV, Bunkóczi G, Chen VB, Davis IW, Echols N, et al. PHENIX: a comprehensive Python-based system for macromolecular structure solution. Acta Crystallogr D Biol Crystallogr. 2010;66(Pt 2):213–21. Epub 20100122. doi: 10.1107/s0907444909052925. PubMed PMID: 20124702; PubMed Central PMCID: PMCPMC2815670.

45. Emsley P, Lohkamp B, Scott WG, Cowtan K. Features and development of Coot. Acta Crystallogr D Biol Crystallogr. 2010;66(Pt 4):486–501. Epub 20100324. doi: 10.1107/s0907444910007493. PubMed PMID: 20383002; PubMed Central PMCID: PMCPMC2852313.

46. Sievers F, Wilm A, Dineen D, Gibson TJ, Karplus K, Li W, et al. Fast, scalable generation of high-quality protein multiple sequence alignments using Clustal Omega. Mol Syst Biol. 2011;7:539. Epub 20111011. doi: 10.1038/msb.2011.75. PubMed PMID: 21988835; PubMed Central PMCID: PMCPMC3261699.

47. Wang EQ, Bukowski JF, Yunis C, Shear CL, Ridker PM, Schwartz PF, et al. Assessing the potential risk of cross-reactivity between anti-bococizumab antibodies and other anti-PCSK9 monoclonal antibodies. BioDrugs. 2019;33(5):571–9. doi: 10.1007/s40259-019-00375-0. PubMed PMID: 31529318; PubMed Central PMCID: PMCPMC6790354.

48. Williams CJ, Headd JJ, Moriarty NW, Prisant MG, Videau LL, Deis LN, et al. MolProbity: More and better reference data for improved all-atom structure validation. Protein Sci. 2018;27(1):293–315. Epub 20171127. doi: 10.1002/pro.3330. PubMed PMID: 29067766; PubMed Central PMCID: PMCPMC5734394.

